# Discovery of novel thrips vector proteins that bind to the viral attachment protein of the plant bunyavirus, tomato spotted wilt virus

**DOI:** 10.1101/416560

**Authors:** Ismael E. Badillo-Vargas, Yuting Chen, Kathleen M. Martin, Dorith Rotenberg, Anna E. Whitfield

## Abstract

The plant-pathogenic virus, tomato spotted wilt virus (TSWV), encodes a structural glycoprotein (G_N_) that, like with other bunyavirus/vector interactions, serves a role in viral attachment and possibly entry into arthropod vector host cells. It is well documented that *Frankliniella occidentalis* is one of seven competent thrips vectors of TSWV transmission to plant hosts, however, the insect molecules that interact with viral proteins, such as G_N_, during infection and dissemination in thrips vector tissues are unknown. The goals of this project were to identify TSWV-interacting proteins (TIPs) that interact directly with TSWV G_N_ and to localize expression of these proteins in relation to virus in thrips tissues of principle importance along the route of dissemination. We report here the identification of six TIPs from first instar larvae (L1), the most acquisition-efficient developmental stage of the thrips vector. Sequence analyses of these TIPs revealed homology to proteins associated with the infection cycle of other vector-borne viruses. Immunolocalization of the TIPs in L1s revealed robust expression in the midgut and salivary glands of *F. occidentalis*, the tissues most important during virus infection, replication and plant-inoculation. The TIPs and G_N_ interactions were validated using protein-protein interaction assays. Two of the thrips proteins, endocuticle structural glycoprotein and cyclophilin, were found to be consistent interactors with G_N_. These newly discovered thrips protein-G_N_ interactions are essential towards better understanding of transmission of persistent propagative plant viruses by their vectors, as well as for developing new strategies of insect pest management and virus resistance in plants.

**Importance Statement:** Thrips-transmitted viruses cause devastating losses to numerous food crops worldwide. For negative-sense RNA viruses that infect plants, the arthropod serves as a host as well by supporting virus replication in specific tissues and organs of the vector. The goal of this work was to identify vector/host proteins that bind directly to the viral attachment protein and thus may play a role in the infection cycle in the insect. Using the model plant bunyavirus, tomato spotted wilt virus (TSWV), and the most efficient thrips vector, we identified and validated six TSWV-interacting proteins from *Frankliniella occidentalis* first instar larvae. Two proteins, an endocuticle structural glycoprotein and cyclophilin, were able to interact directly with the TSWV attachment protein, G_N_, in insect cells. The TSWV G_N_-interacting proteins provide new targets for disrupting the virus-vector interaction and could be putative determinants of vector competence.

## Introduction

Vector-borne diseases caused by animal- and plant-infecting viruses are some of the most important medical, veterinary, and agricultural problems worldwide (1, 2). The majority of viruses infecting plants and animals are transmitted by arthropods. Understanding the viral and arthropod determinants of vector competence is important for basic knowledge of virus-vector interactions and development of new interdiction strategies to control disease. Significant progress has been made towards identification of viral determinants of transmission, but the interacting molecules in vectors remain largely elusive. For negative-sense RNA viruses, vector factors that mediate the transmission process have not been well characterized.

*Bunyavirales* is the largest order of negative-sense RNA viruses; twelve families are described (http://www.ictvonline.org/virustaxonomy.asp). The *Bunyavirales* contains plant and insect vector-infecting viruses that make up the Family *Tospoviridae* (3–5). Within this family, there are eighteen species and several unassigned viruses that most likely will be classified as unequivocal members of the *Orthotospovirus* genus. *Tomato spotted wilt orthotospovirus* is the type species within this genus and has been best characterized in terms of viral host range, genome organization and protein functions (6, 7).

Tomato spotted wilt virus (TSWV) infects both monocotyledonous and dicotyledonous plants encompassing more than 1,000 plant species worldwide (8). Due to the extremely wide host range, TSWV has caused severe economic losses to various agricultural, vegetable and ornamental crops. The TSWV virion has a double-layered, host-derived membrane studded with two glycoproteins (G_N_ and G_C_) on the surface. The viral glycoproteins play an essential role in attachment to the thrips gut and fusion of the virus and host membrane (7, 9–11). Virus particles range in size from 80 to 120 nm in diameter, and inside the particle are three genomic RNAs designated long (L), medium (M) and small (S) RNA based on the relative size of each molecule.

Although TSWV can be maintained in the laboratory through mechanical inoculation, it is transmitted in nature by insect vectors commonly known as thrips (Order Thysanoptera, Family *Thripidae*). Five species of *Frankliniella* and two species of *Thrips* are reported to be the vectors of TSWV (6). Among these species, the western flower thrips, *Frankliniella occidentalis* Pergande, is the most efficient vector of TSWV and it has a worldwide distribution. TSWV is transmitted by thrips vectors in a persistent propagative manner, and the midgut cells and primary salivary glands are two major tissues for TSWV replication (12, 13). Only thrips that acquire virus during the early larval stage are inoculative as adults (13–15). Because the TSWV G_N_ protein has been identified to bind to thrips midguts and play a role in virus acquisition by thrips (9–11), we sought to identify thrips proteins that interact directly with G_N_, the viral attachment protein (16). Using gel overlay assays to identify first instar larval (L1) proteins that bind to purified virions or G_N_, we discovered six TSWV-interacting proteins (TIPs) from *F. occidentalis*. Identification of these proteins using mass spectrometry was followed with secondary assays to validate the interactions and characterize protein expression in larval thrips. Two TIPs, an endocuticle structural glycoprotein and cyclophilin, interacted with G_N_ and co-localized with G_N_ when co-expressed in insect cells. These thrips proteins may play a role in virus entry or mediate other steps in the virus infection process in thrips. These proteins represent the first thrips proteins that bind to TSWV proteins, and these discoveries provide insights toward a better understanding of the molecular interplay between vector and virus.

## Results

### Identification of bound *F. occidentalis* larval proteins using overlay assays

Proteins extracted from first instar larvae bodies were separated by 2-D electrophoresis, and overlay assays were performed with purified TSWV virions or recombinant G_N_ glycoprotein to identify bound thrips proteins. Virion overlays identified a total of eight proteins spots (Fig. 1) - three occurred consistently in all four biological replicates, while five were present in three. Mass spectrometry and subsequent peptide sequence analysis against a 454-transcriptome database (*Fo* Seq) identified one to four different transcript matches per spot (Table 1), where in four cases, the same putative transcript matched peptides in more than one spot. Using recombinant G_N_ glycoprotein, 11 protein spots were detected in both biological replicates of the overlay assay (Fig. 2), and each spot was comprised of a single protein (single transcript match) occurring in multiple spots - there were a total of two different G_N_-interacting proteins represented by the 11 spots (Table 2). For each overlay experiment that was run, a control blot was included to identify background, *i. e*., non-specific binding by the primary and secondary antibodies, demonstrating detection of the positively identified spots well-exceeded background (Fig. 1 and 2). In an additional gel overlay assay using virus-free plant extract (mock purification) obtained from healthy *D. stramonium* plants, no protein spots above the antibody control were detected (data not shown).

**Fig 1.**
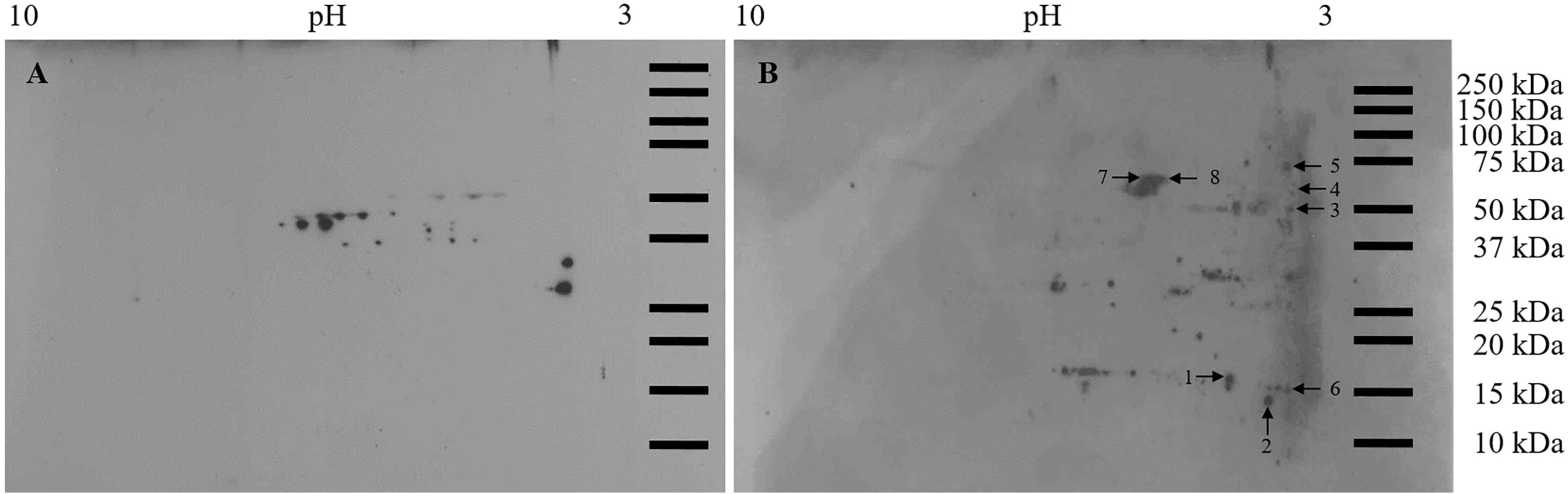
Overlay assay using purified virions and *F. occidentalis* first instar proteins resolved in two-dimensional gels. Total proteins (150 μg) extracted from pooled healthy first instar larvae (0-17-hour old) of *F. occidentalis* were resolved by 2-D gel electrophoresis and transferred to nitrocellulose membranes. After blocking, membranes were incubated overnight with (A) blocking buffer (negative control) or (B) purified TSWV at 25 μg/mL, and then incubated with polyclonal rabbit anti-TSWV G_N_ antiserum. Only protein spots that consistently bound to purified TSWV in three (spots 1, 2, 4, 6, and 7) and four (spots 3, 5, and 8) biological replicates of the overlay assay were collected from three individual picking gels and subjected to ESI mass spectrometry for protein identification. Protein spots observed in the no-overlay-control membrane represent non-specific binding and were not collected for further analysis. Molecular mass (in kilodaltons) is shown on the Y axis and pI (as pH range) is shown on the X axis.

**Fig 2.**
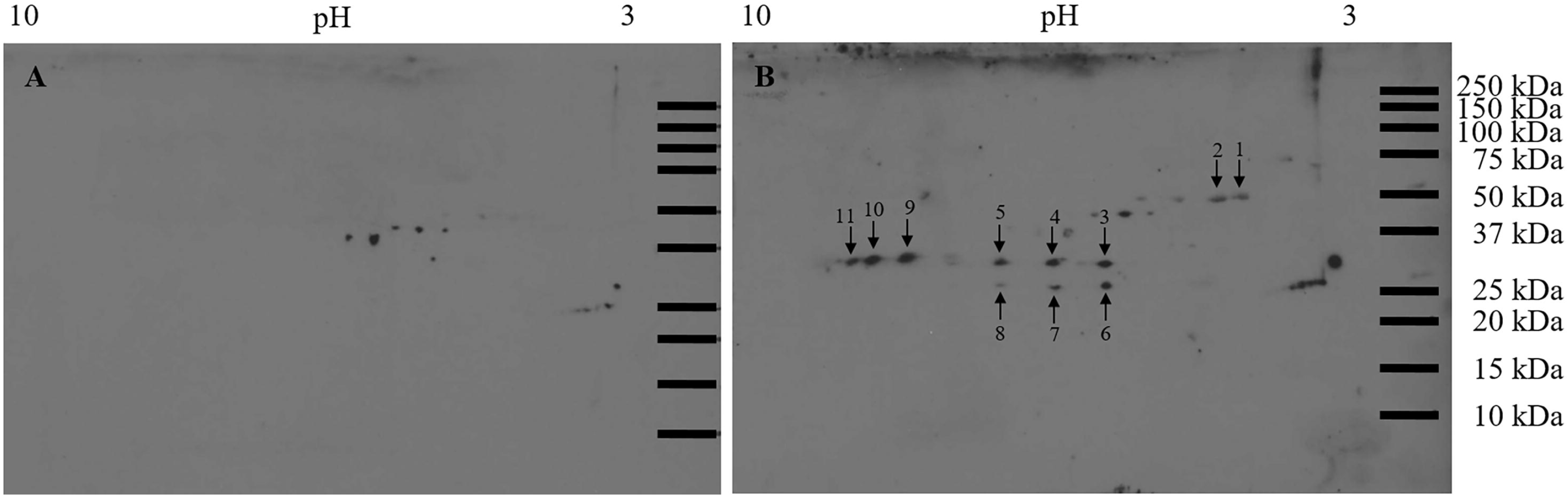
Overlay assay using recombinant G_N_ and *F. occidentalis* first instar proteins resolved in two-dimensional gels. 150 μg total proteins extracted from pooled healthy first instar larvae (0-17-hour old) of *F. occidentalis* were resolved by 2-D gel electrophoresis and transferred to nitrocellulose membranes. The membranes were incubated overnight with (A) blocking buffer (negative control) or (B) recombinant TSWV G_N_ (3.5 μg/mL). Using the polyclonal rabbit anti-TSWV G_N_, protein spots that consistently bound to the recombinant TSWV G_N_ in two (spots 1 through 11) biological replicates of the overlay assay were collected from two individual picking gels and subjected to ESI mass spectrometry for protein identification. Protein spots observed in the no-overlay-control membrane represent non-specific binding and were not collected for further analysis. Molecular mass (in kilodaltons) is shown on the Y axis and pI (as pH range) is shown on the X axis.

**Table 1.**
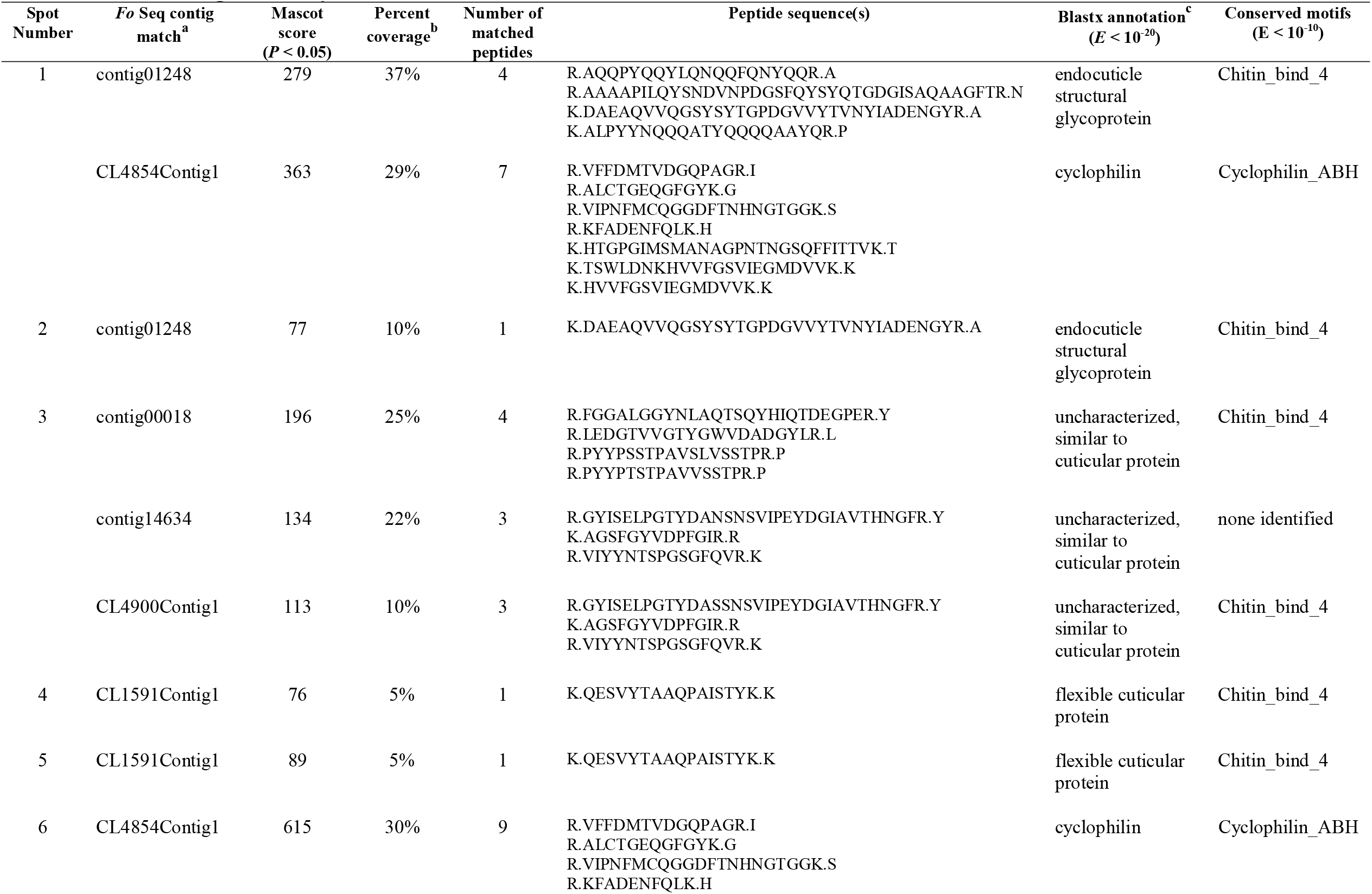
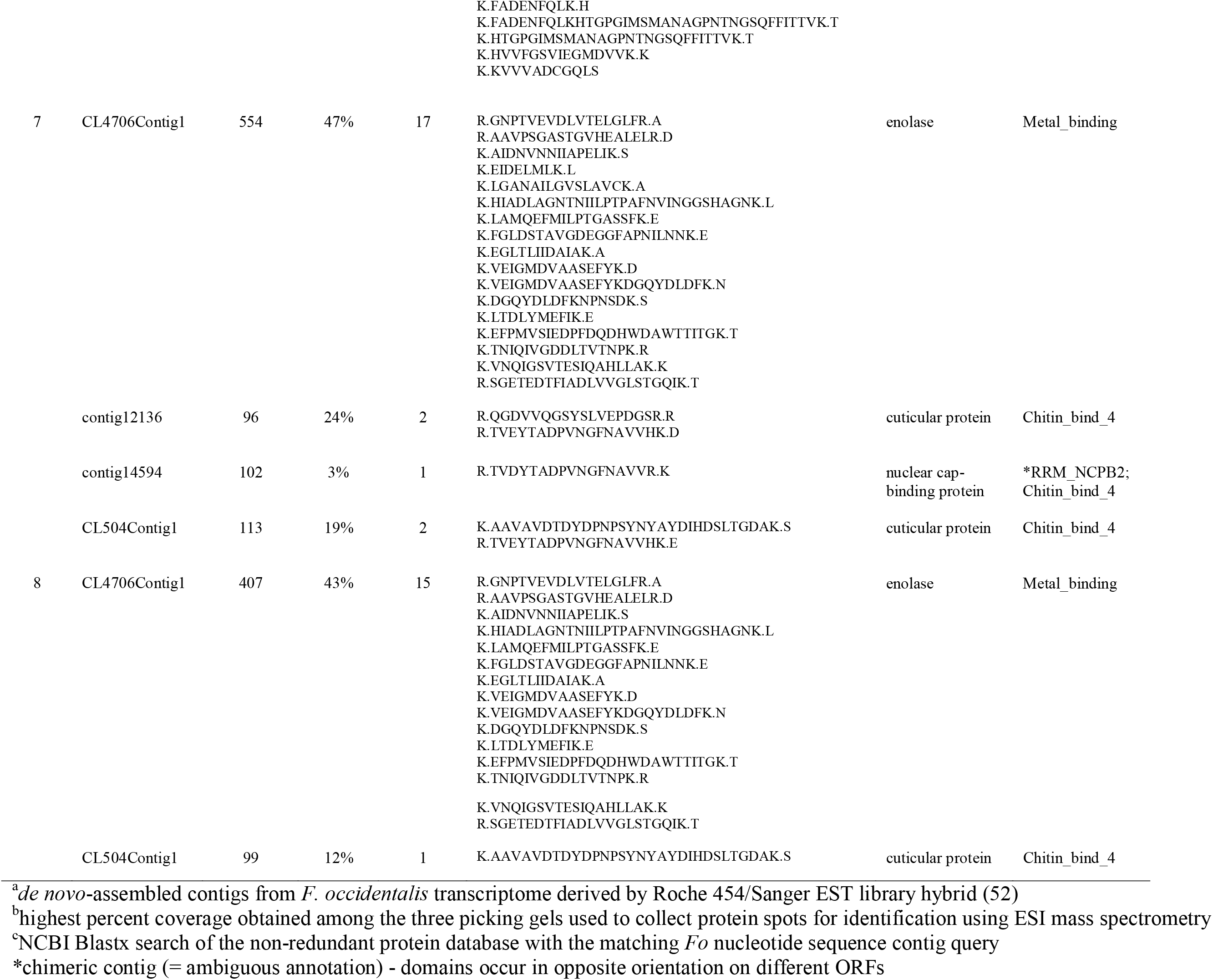
Identification of *Frankliniella occidentalis* larval proteins bound to purified virions of *Tomato spotted wilt virus* in twodimensional (2-D) gel overlays.

### Annotation of six candidate TSWV-interacting proteins (TIPs)

Our stringent sequence-filtering criteria retained four different virion-interacting proteins [endocuticle structural glycoprotein: endoCP-V (contig01248, GenBank accession: MH884756); cuticular protein: CP-V (CL4900Contig1, MH884758), cyclophilin (CL4854Contig1, MH884760), and enolase (CL4706Contig1, MH884759), Table 1] and two G_N_-interacting proteins [mitochondrial ATP synthase α, mATPase (CL4310Contig1, MH884761) and endocuticle structural glycoprotein; endoCP-G_N_ (CL4382Contig1, MH884757), Table 2] to move forward to validation and biological characterization. Collectively, these six protein candidates are referred to as ‘<u>T</u>SWV-<u>I</u>nteracting <u>P</u>roteins’ or TIPs and their putative identifications and sequence features are shown in Table 3. Blastp analysis of the predicted, longest complete ORFs confirmed their annotations and putative sequence homology to proteins in other insects. The three cuticle-associated TIPs (endoCP-G_N_, endoCP-V, and CP-V) contained predicted signal peptide sequences, indication of secreted proteins, and a chitin-binding domain (CHB4). Pairwise alignments (Blastp) between the translated ORFs of the three cuticle TIPs and the six other gel overlay-resolved CPs or endoCPs revealed sequence diversity; where matches among the different cuticle proteins occurred (cut-off = *E* < 10^−3^), % amino acid identities ranged from 53% – 67%, covering 30% – 49% of the queries, with e-values ranging from 2.4 × 10^−2^ – 3.6 × 10^−24^. The only exception was the CP-V and contig00018 alignment, which appeared to be 100% identical along the entire length of contig00018 (*E* = 2.6 × 10^−162^) (data not shown). The other three TIPs (cyclophilin, enolase and mATPase) contained motifs characteristic of these proteins (Table 3).

**Table 2.**
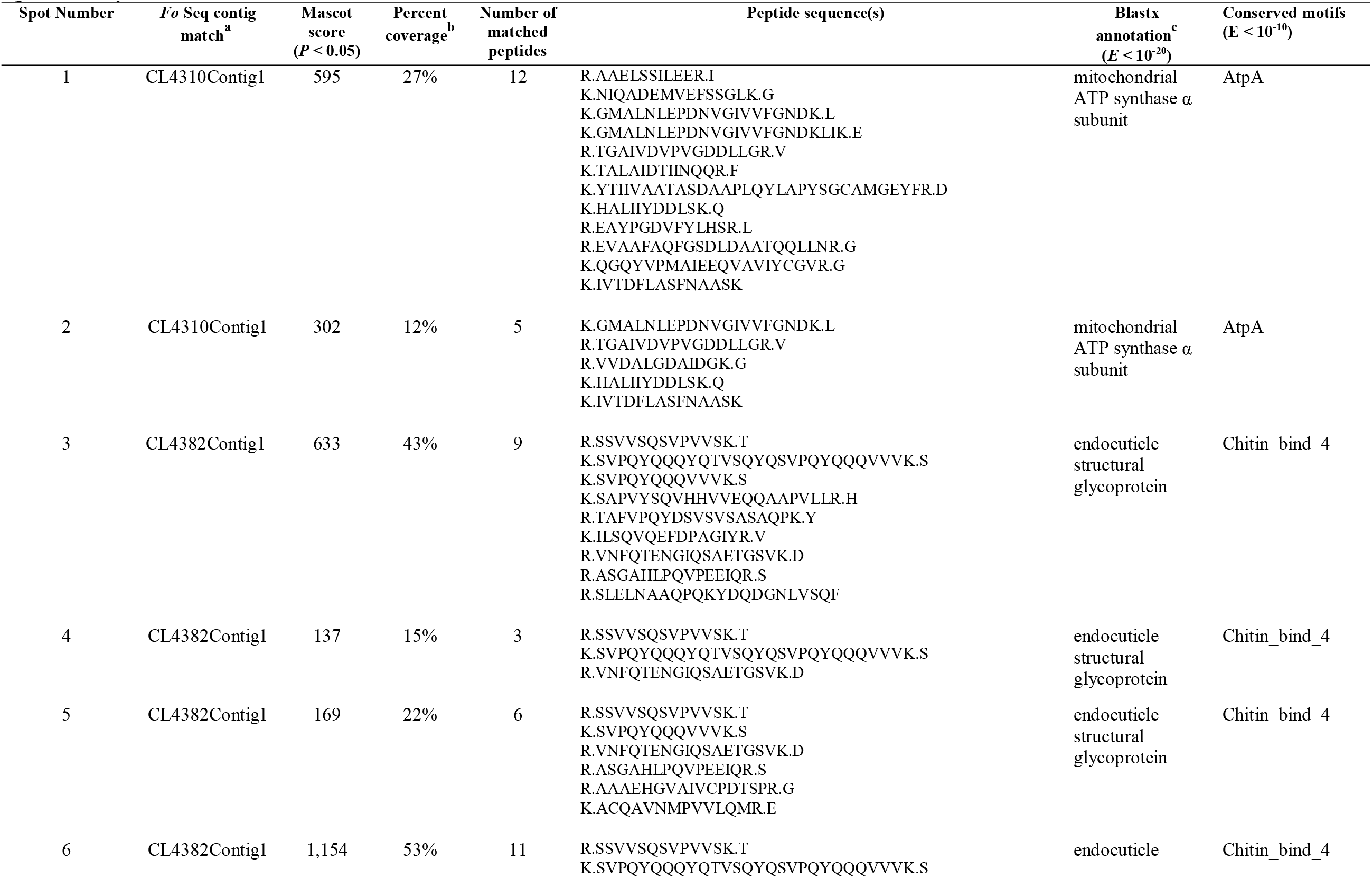
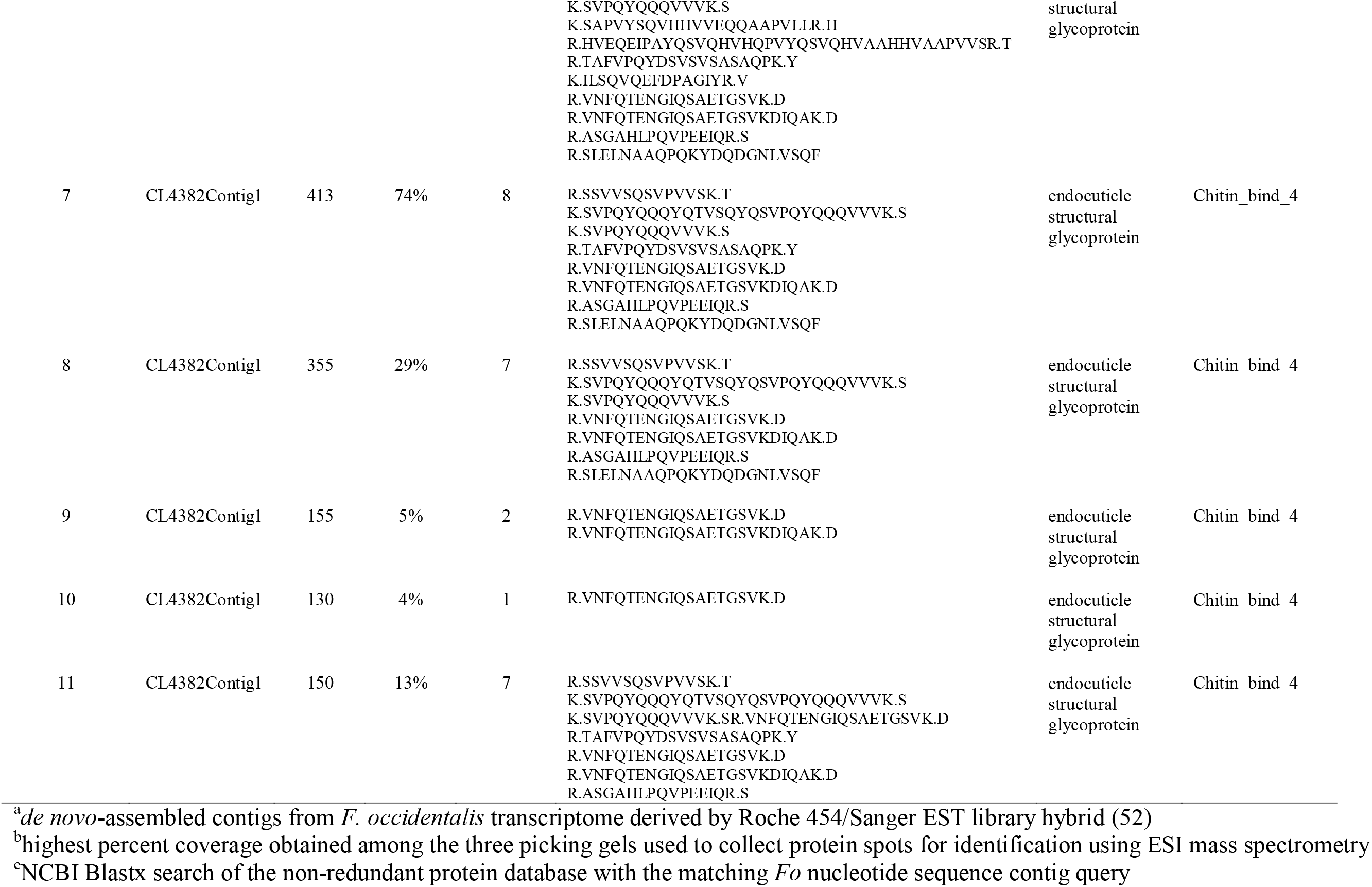
Identification of *Frankliniella occidentalis* larval proteins bound to recombinant glycoprotein-N (G_N_) in two-dimensional (2-D) gel overlays.

**Table 3.** Final candidate list of six TSWV-interacting proteins (TIPs)^a^ from larval *Frankliniella occidentalis* to move forward to validation and biological characterization.

| Putative protein | Fo Seq contig match<sup>b</sup> | ORF length<sup>c</sup><br>(nt/aa) | Signal peptide<sup>d</sup><br>(aa position) | Conserved domains<sup>e</sup> |  |  | Top Genbank match<sup>f</sup><br>(% coverage, % identity, *E*) |
| --- | --- | --- | --- | --- | --- | --- | --- |
|  |  |  |  | Name | Position<br>(aa) | E-value |  |
| Cuticular protein-V (CP-V) | CL4900Contig1 | 1,302/434 | 1-18 | Chitin_bind_4 | 447-92 | 5.7x10<sup>-6</sup> | XP_017786818.1: PREDICTED: cell surface glycoprotein 1 [*Nicrophorus vespilloides*] (94%, 44%, 1x10<sup>-85</sup>) |
| Endocuticle structural glycoprotein-G<sub>N</sub> (endoCP-G<sub>N</sub>) | CL4382Contig1 | 852/283 | 1-15 | Chitin_bind_4 | 190-246 | 7.5x10<sup>-11</sup> | XP_018334183.1: endocuticle structural glycoprotein SgAbd-2-like [*Agrilus planipennis*] (33%, 53%, 4x10<sup>-22</sup>) |
| Endocuticle structural glycoprotein-V (endoCP-V) | contig01248 | 522/173 | 1-17 | Chitin_bind_4 | 63-119 | 1.4x10<sup>-18</sup> | XP_022906571.1: endocuticle structural glycoprotein SgAbd-2-like [*Onthophagus taurus*] (63%, 58%, 2x10<sup>-35</sup>) |
| Cyclophilin (peptidyl-prolyl cis-trans isomerase) | CL4854Contig1 | 618/205 | None | cyclophilin_ABH_like<br>PpiB<br>Pro_isomerase | 44-202<br>57-196<br>47-201 | 2.3x10<sup>-120</sup><br>2.6x10<sup>-63</sup><br>1.2x10<sup>-62</sup> | XP_019753975.1: PREDICTED: peptidyl-prolyl cis-trans isomerase [*Dendroctonus ponderosae*] (99%, 75%, 2x10<sup>-109</sup>) |
| Enolase | CL4706Contig1 | 1,302/433 | None | PLN00191<br>enolase<br>Eno<br>Enolase_C | 3-433<br>5-417<br>6-431<br>143-433 | 0<br>0<br>0<br>0 | XP_019767728.1: PREDICTED: enolase-like [*Dendroctonus ponderosae*] (99%, 87%, 0) |
| Mitochondrial ATP synthase $\alpha$ | CL4310Contig1 | 1,665/554 | None | PRK09281<br>AtpA<br>F1_ATPase_alpha<br>ATP-synt_ab | 45-551<br>45-553<br>135-416<br>190-413 | 0<br>0<br>0<br>6.1x10<sup>-117</sup> | XP_023718907.1: ATP synthase subunit alpha, mitochondrial [*Cryptotermes secundus*] (99%, 89%, 0) |

<sup>a</sup>sequences deposited into National Center for Biotechnology Information (NCBI) GenBank with the following accessions: CP-V (MH884758), endoCP-G<sub>N</sub> (MH884757), endoCP-V (MH884756), cyclophilin (MH884760), enolase (MH884759), and mitochondrial ATP synthase $\alpha$ (MH884761).
<sup>b</sup>*de novo*-assembled contigs from *F. occidentalis* transcriptome derived by Roche 454/Sanger EST library hybrid (52)
<sup>c</sup>prediction by NCBI ORF Finder (https://www.ncbi.nlm.nih.gov/orffinder/)
<sup>d</sup>prediction by Signal P (http://www.cbs.dtu.dk/services/SignalP/)
<sup>e</sup>prediction by NCBI Batch Web CD Search Tool (https://www.ncbi.nlm.nih.gov/Structure/bwrpsb/bwrpsb.cgi) (CDSEARCH/cdd v3.16); E-value cut-off = $10^{-5}$ , only specific hits shown
<sup>f</sup>Blastp search of NCBI nr protein database against TIP ORF as of August 06, 2018; top match indicates highest alignment (max) score

### Classification of cuticular TIPs

All three cuticular TIPs were classified as members of the Cuticle Protein - R&R Consensus motif (CPR) family (17) based on the occurrence of one RR extended consensus CHB4, with both endoCP-G_N_ (*E* = 4 × 10^−18^) and endoCP-V (*E* = 1 × 10^−26^) predicted to belong to the RR1 group, and CP-V weakly supported (*E* = 5×10^−6^) to belong to the RR2 group of CPRs. All three sequences were phylogenetically placed into the RR1 major clade with strong bootstrap support (82%, Fig. S1) in relation to other *F. occidentalis* CPRs previously found to be downregulated in TSWV-infected first instar larvae (18) and CPRs of other insect species. Within the RR1 clade, the CP-V CHB4 domain clustered with a CP of the small brown planthopper, *Laodelphax striatella* (KC485263.1, CprBJ), reported to bind to the nucleocapsid protein pc3 of rice stripe virus (RSV) during infection of the vector (19) and which was predicted (*E* = 5 × 10^−7^) to be classified in the RR1 group.

### Antisera show specificity against each TIP-peptide

The antisera specifically bound to their TIP peptides in dot-blot assays (Fig. 3), although the affinity of each antibody to its cognate TIP peptide varied. The mATPase antibody had highest affinity to mATPase peptide, while the CP-V antibody had lowest affinity to CP-V peptide (the high concentration of CP-V peptide, 2.5 mg/mL was used, and all primary and secondary antibody incubation time was doubled, and the developing time for chemiluminescence detection was increased). This result demonstrates the specificity of the TIP-peptide antibodies that were used in subsequent localization experiments with first instar thrips larvae.

**Fig. 3.**
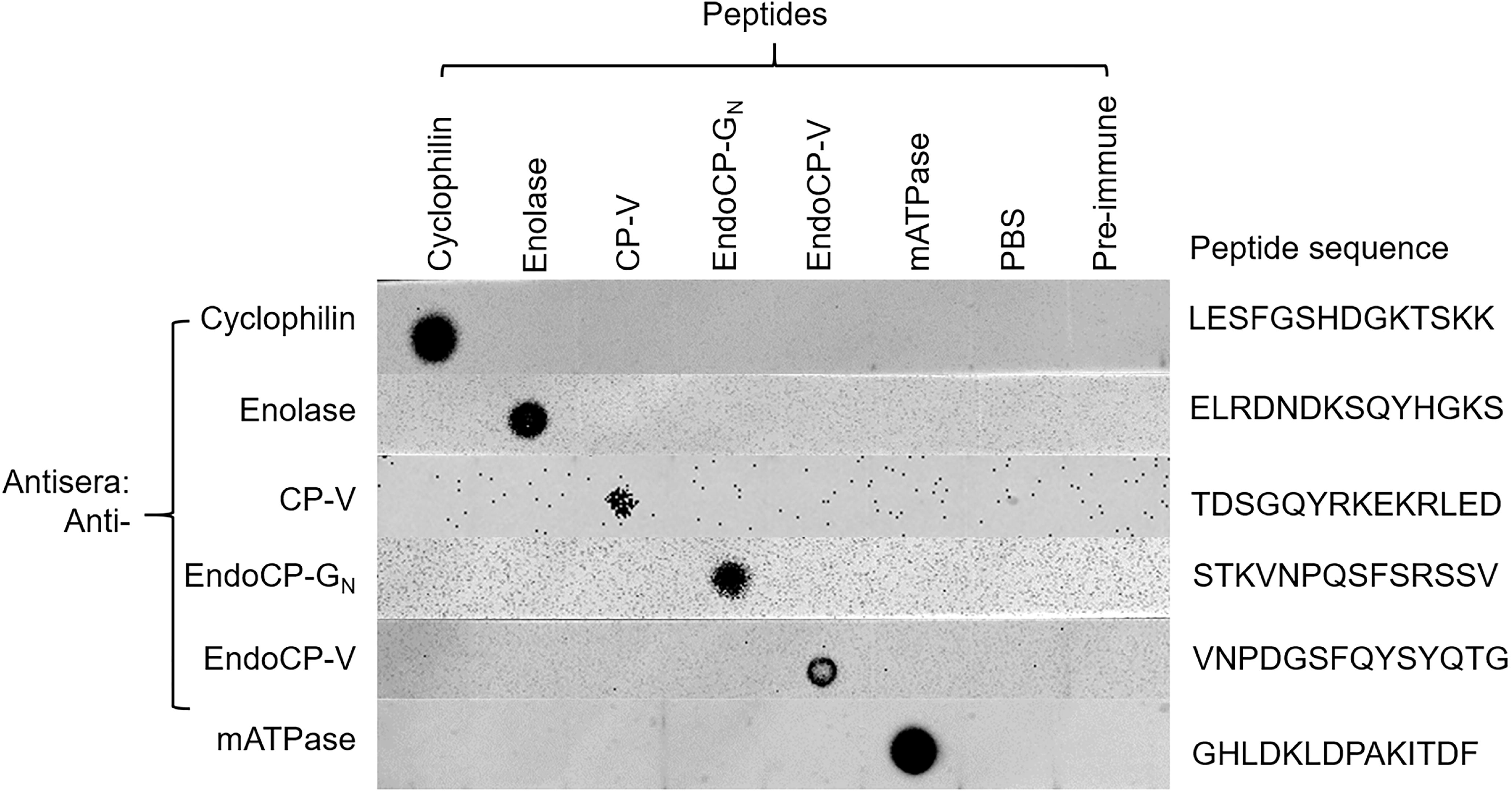
Antisera specificity against each TSWV-interacting protein (TIP) (peptide) using dot blot analysis. Peptides that were used for producing its antibody were diluted to 100 μg/mL (for cyclophilin, enolase, endoCP-G_N_, endoCP-V, and mATPase), and 2.5 mg/mL (for CP-V), and 2ul of each peptide were used for each test. PBS buffer and pre-immune serum (500,000 × dilution) were used as controls. All six diluted peptides and two controls were loaded onto six nitrocellulose membrane strips. Each strip was first incubated with one specific primary antibody (0.5 μg/mL, generated in mice), then incubated with goat anti-mouse-HRP (1:5,000 dilution). Each membrane strip was developed independently.

### *In vivo* localization of TIPs in *F. occidentalis* in midguts and salivary glands

Specific antisera raised against each confirmed TIP was used in immunolabeling experiments to localize protein expression in L1 tissues *in vivo*. Visualization by confocal microscopy revealed that all six TIPs were primarily localized at the foregut (esophagus), midgut (epithelial cells and visceral muscle), salivary glands (including both primary and tubular salivary glands), and Malpighian tubules (Fig. 4), and this was the case in 100% of the dissected tissues treated with TIP-specific antisera. It was difficult to completely dissect and separate hindgut from the carcass without damaging the tissue, therefore, the localization of TIPs in the hindgut was unclear. For each experimental replicate and unique antibody, controls of secondary antibody only and pre-immune serum plus secondary antibody were conducted and visualized by confocal microscopy. The confocal laser settings (power and percent/gain) were adjusted to remove any background fluorescence observed with pre-immune serum for each TIP as they showed slightly higher background compared to the secondary antibody control. The bright field and merged images of these controls, depicting actin- and nuclei-labeling, are shown in Fig. S2.

**Fig 4.**
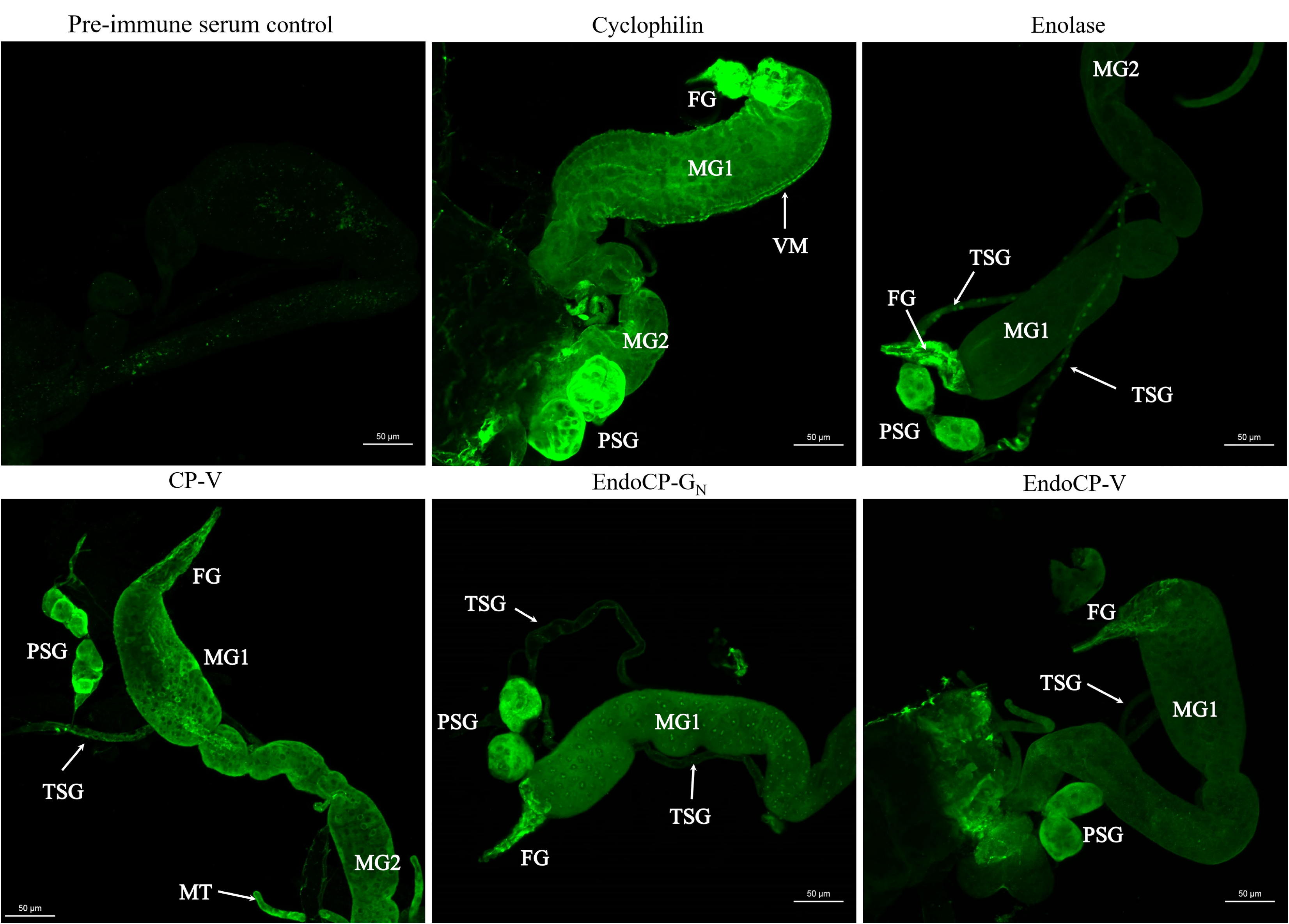
Immunolabeling of TSWV-interacting proteins (TIPs) within first instar larvae of *F. occidentalis*. The synchronized first instar larvae (0-17-hour old) were kept on 7% sucrose solution for 3 hours to clean their guts from plant tissues. These larvae were then dissected and immunolabeled using specific antibodies against each TIP as indicated. Thrips tissues incubated with pre-immune mouse serum are depicted here. Confocal microscopy detection of green fluorescence (Alexa Fluor 488) represents the localization of each TIP. TIPs were mainly localized at foregut (FG), midgut (MG), which includes epithelial cells and visceral muscle (VM), principle salivary glands (PSG), tubular salivary glands (TSG), and Malpighian tubules (MT). All scale bars are equal to 50 μm.

### Validation of interactions between TIPs and TSWV G_N_ using BiFC

Before launching a BiFC analysis of candidate protein interactions *in planta*, it is critical to determine if position of a fused fluorescent protein tag (N- or C-terminus of the candidate protein) affects the expression and/or localization of the fusion protein in cells. Furthermore, it was expected that the signal peptides located on the N-terminus of the soluble (G_N_-S) and insoluble (G_N_) TSWV glycoprotein, and the cuticular TIPs (CP-V, endoCP-V, endoCP-G_N_), would preclude placement of tags at the N-terminus of these proteins. GFP fused to the N-terminus of the glycoprotein (G_N_ and G_N_-S), the cuticle TIPs (endoCP-G_N_ and endoCP-V), and mATPase α produced weak signal or reduced mobility in the cell (data not shown). For the remaining proteins, there was no effect of fluorescent-protein tag location on protein expression or mobility. Thus, all protein localization and BiFC validation experiments were performed with C-terminally fused TIPs for consistency in the assays.

The GFP-TIP fusions displayed distinct cellular localization patterns when expressed in plants (Fig. S3). Cyclophilin and mATPase appeared to be localized to the nuclei and along the cell periphery, while enolase and CP-V were present in the membranes surrounding the nuclei as well as the cell periphery. Both endoCP-G_N_ and endoCP-V had a punctate appearance outside of the nucleus. All three cuticular TIPs (CP-V, endoCP-G_N_, endoCP-V) formed small bodies that appeared to be moving along the endo-membranes of the cell, consistent with secretion. All TIPs were co-localized with the ER marker; however, none appeared to be co-localized with the Golgi marker (data not shown).

BiFC analysis validated the TSWV-TIPs interactions identified in the overlay assays between virions and G_N_ and enolase, m-ATPase, endoCP-G_N_ and endoCP-V (Fig. 5). We used the soluble form of the viral glycoprotein (G_N_-S) and the insoluble form with the transmembrane domain and cytoplasmic tail in BiFC assays. The insoluble form of G_N_ interacted with enolase, endoCP-G_N_ and endoCP-V. The proposed ectodomain of G_N_-S interacted with mATPase and endoCP-V. All of the BiFC interactions were detected in the membranes surrounding the nuclei and at the cell periphery, generally consistent with the localization patterns of the GFP-fused TIPs as described for the localization experiment above (Fig. S3).

**Fig 5.**
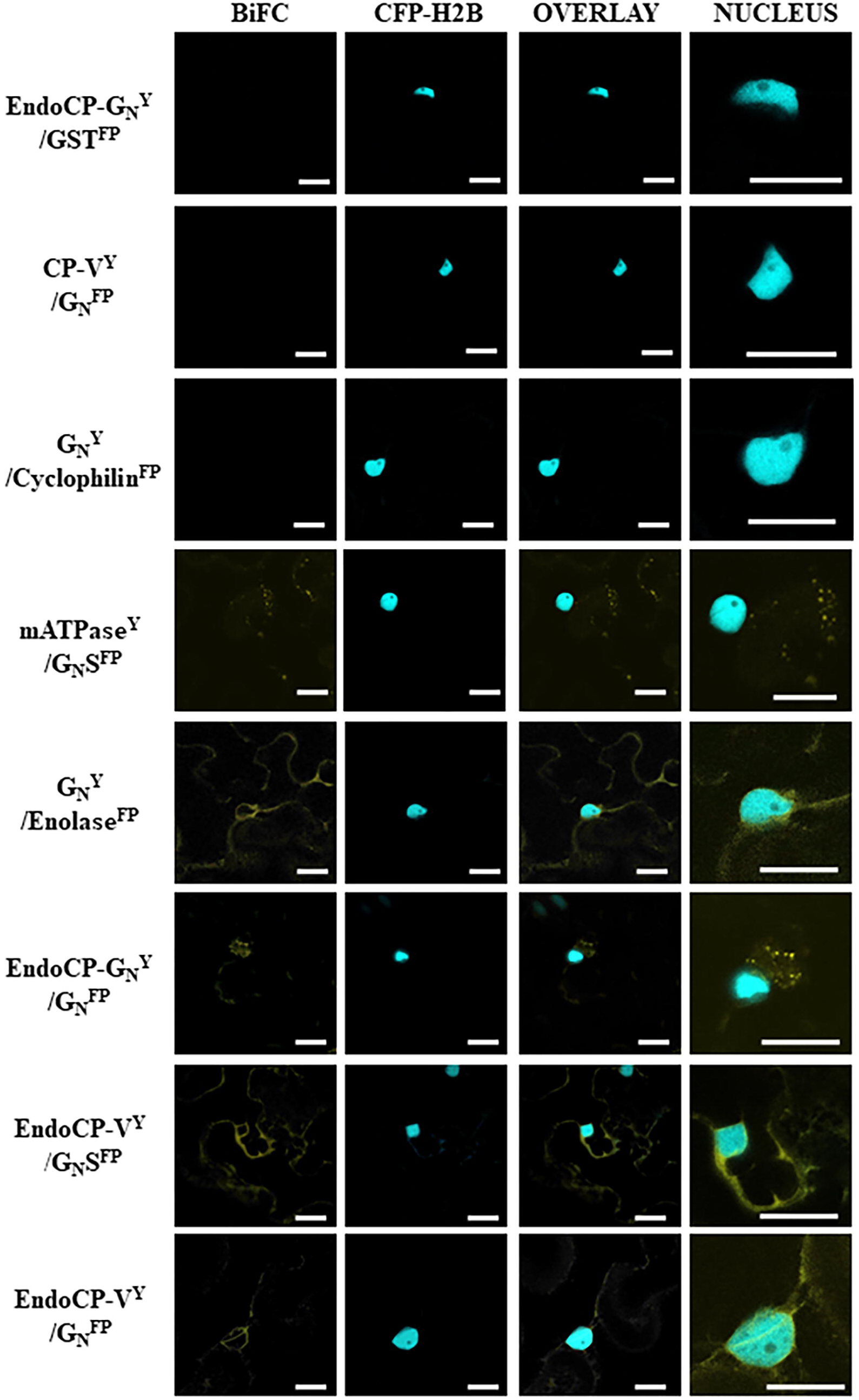
Confirmation of interactions between TSWV proteins and TSWV-interacting proteins (TIPs) using bimolecular fluorescence complementation (BiFC) in *Nicotiana benthamiana*. Plants transgenic for a nuclear marker fused to cyan fluorescent protein (CFP-H2B) were infiltrated with suspensions of *Agrobacterium tumefaciens* transformed with plasmids encoding the G_N_ protein (full length or soluble form, G_N_-S) and TIPs proteins (EndoCP-G_N_, CP-V, Cyclophilin, mATPase, Enolase, EndoCp-V) fused to either the amino or carboxy terminus of yellow fluorescent protein (YFP). The designation of Y indicates this is the n-terminal half of YFP and FP represents the c-terminal half of YFP. The Y or FP position in the name indicates all are carboxy terminal fusions to the protein of interest. The positive interactors are seen by fluorescence of YFP in images shown in the ‘BiFC’ column. The ‘CFP-H2B’ column is indicated to give cellular reference, and the overlay between the two is also shown. The final column is the nucleus enlarged to show detail of the interacting TIPs within the cellular context. The first row is a representative negative control with a TIP and glutathione S-transferase (all thrips and virus proteins were tested with the negative control to rule out non-specific interactions). All scale bars are equal to 20 μm.

### Validation of gel overlay protein-protein interactions using the split-ubiquitin membrane-based yeast two-hybrid analysis

The split-ubiquitin membrane-based yeast two-hybrid (MbY2H) system was used to validate the gel overlay interactions between the six candidate TIPs and TSWV glycoprotein G_N_. The presence of a transmembrane domain near the C-terminus of TSWV G_N_ makes the MbY2H system the best choice for validation of TSWV G_N_ interactions with the candidate TIPs. The interaction between G_N_ and endoCP-G_N_ was consistent and strong based on the number of colonies growing on QDO, *i.e*., - more than 1,000 colonies on all QDO plates for all three replicates (Fig 6A), and this interaction was confirmed by β-galactosidase assay (Table S1). We detected a consistent but weak interaction (average of 15 colonies) between G_N_ and cyclophilin, and seven of nine colonies tested by β-galactosidase assay were positive. The remaining four TIPs showed no interaction with G_N_ using MbY2H. Contrary to the MbY2H results, G_N_ was determined to interact with enolase and endoCP-V in BiFC experiments. The steric constraints imposed by the position (C- or N-terminus) of the reporter in the MbY2H (Ubiquitin half) and BiFC (YFP half) systems in yeast versus plants cells, respectively, may explain the contrasting interactions observed in these assays.

**Fig 6.**
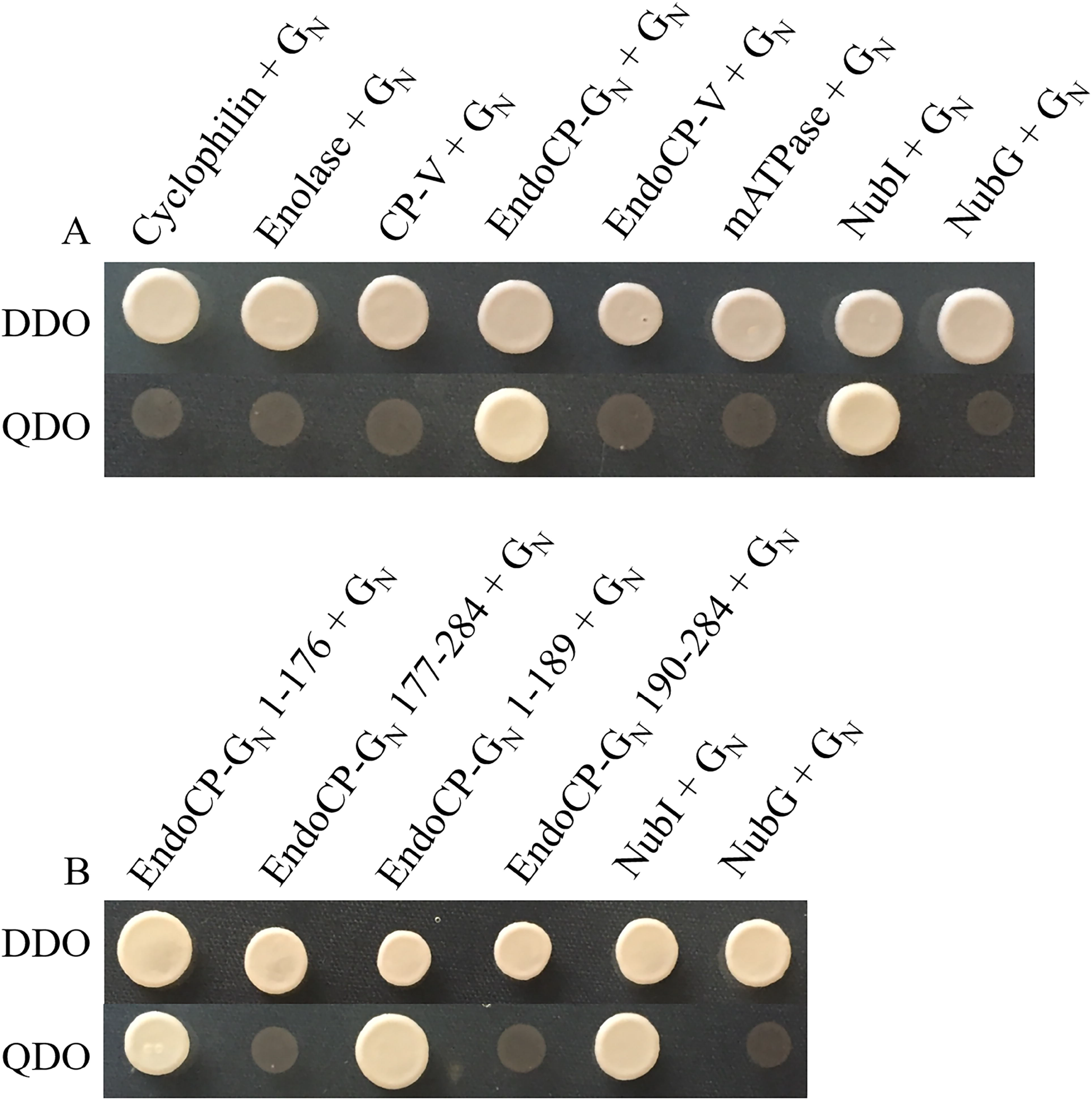
Validation of TSWV-interacting proteins (TIPs) with G_N_, and identification of the interacting domain of endoCP-G_N_ using split-ubiquitin membrane-based yeast two hybrid (MbY2H). (A) Interactions between G_N_ and six TIPs. G_N_ was expressed as G_N_-Cub, and TIPs were expressed as NubG-TIPs using MbY2H vectors. (B) Interactions between TSWV G_N_ and different regions of endoCP-G_N_. EndoCP-G_N_ was expressed as either the N-terminal domain (amino acids 1-176 and 1-189) that includes the non-conserved region or the C-terminal region (amino acids 177-284 and 190-284) that includes the conserved Chitin_bind_4 motif (CHB4) of endoCP-G_N_. Interactions between G_N_-Cub and NubI, G_N_-Cub and NubG were used as positive and negative controls respectively for all MbY2H assays. Co-transformation of pTSU2-APP and pNubG-Fe65 into NYM51 was used as another positive control (data not shown); DDO = yeast double dropout (SD/-Leu/-Trp) media, and QDO = yeast quadruple dropout (SD/-Ade/-His/-Leu/-Trp) media.

### The non-conserved region of endoCP-G_N_ binds TSWV G_N_

Given the role of G_N_ as the viral attachment protein in the larval thrips midgut epithelium (7, 10) and the confirmed direct interaction between endoCP-G_N_ and TSWV G_N_, there was interest in broadly identifying the amino acid region in the endoCP-G_N_ sequence that binds G_N_. We hypothesized that the non-conserved region of endoCP-G_N_ (N-terminal region up to 176 aa or 189 aa) and not the CHB4 motif might play an important role in the interaction with TSWV G_N_. Using the MbY2H system, it was determined that the non-conserved region of the endoCP-G_N_ sequence had as strong of an interaction with TSWV G_N_ as the complete endoCP-G_N_ sequence (Fig 6B and Table S1) - more than 500 colonies on each QDO plate for each experimental replicate – while the predicted CHB4 motif alone (amino acid positions 190-284) or CHB4 plus few amino acids upstream (position 177-284) did not show an interaction. The non-conserved endoCP-G_N_ sequence region was determined to have no significant matches to sequences in NCBI non-redundant nucleotide and protein databases.

### Cyclophilin and endoCP-G_N_ co-localized with TSWV G_N_ in insect cells

To further explore the interactions between TSWV G_N_ and the two robust thrips interacting proteins, cyclophilin and endoCP-G_N_, we co-expressed the proteins as fusions with GFP or RFP in insect cells. The fusion proteins cyclophilin-RFP and endoCP-G_N_-RFP and TSWV G_N_-GFP were expressed individually and together in Sf9 cells. When fusion proteins cyclophilin-RFP and endoCP-G_N_-RFP were individually expressed in Sf9 cells, they were localized within the entire cytoplasm (Fig. 7). Similarly, the fusion protein TSWV-G_N_-GFP was also expressed in the cytoplasm, but specifically localized at structures that may be ER and/or Golgi, consistent with previous work localizing G_N_ to these organelles in animal cells (20). When cyclophilin-RFP and TSWV-G_N_-GFP or endoCP-G_N_-RFP and TSWV-G_N_-GFP were co-expressed in Sf9 cells, they co-localized within small punctate structures, which was different from their original localization (Fig. 7). However, the controls of co-expressed RFP and GFP (co-transfection of pHRW and pHGW) were distributed throughout the cytoplasm, and the localization of cyclophilin-RFP, endoCP-G_N_-RFP and TSWV G_N_-GFP did not change with the presence of GFP or RFP (Fig. S4). The controls of co-expressed RFP and GFP (co-transfection of pHRW and pHGW) were distributed throughout the cytoplasm in single and double transfections (Fig. 7). Although these unknown co-localization sites need to be further characterized, these co-localization results strongly supports the validity of *in vivo* interactions of cyclophilin and endoCP-G_N_ with TSWV-G_N_.

**Figure 7.**
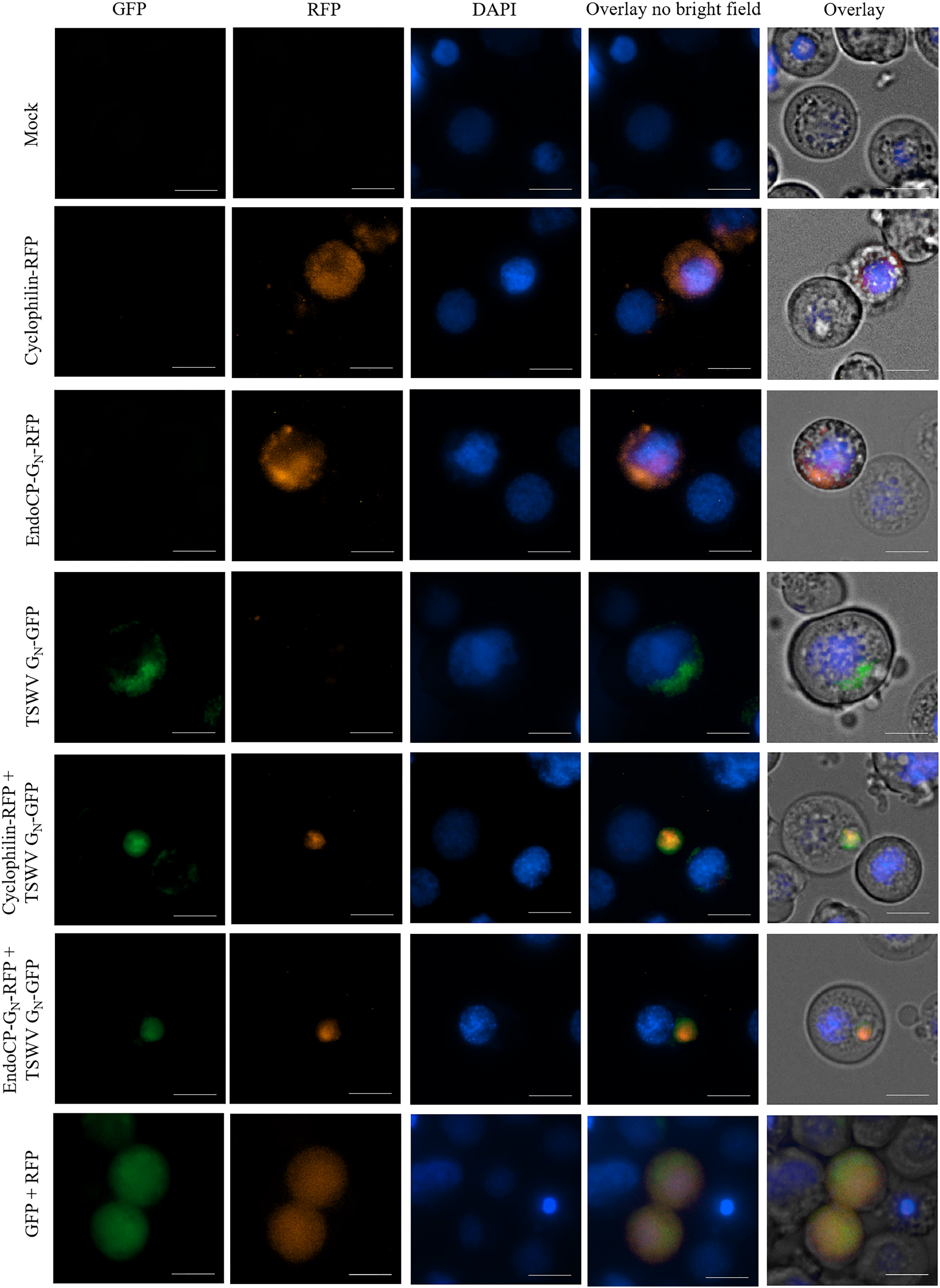
Co-localization of TSWV G_N_ and endoCP-G_N_ or cyclophilin in insect cells. Open reading frames of cyclophilin, endoCP-G_N_, and TSWV G_N_ were cloned into *Drosophila* gateway vectors (with Hsp70 promotor and gateway cloning cassette), and the pHWR and pHWG expression plasmids were used for the following fusion proteins: cyclophilin-RFP, endoCP-G_N_-RFP and TSWV G_N_-GFP. The recombinant plasmids, pHWR-cyclophilin, pHWR-endoCP-G_N_ and pHWG-TSWV G_N_ were single or co-transfected into insect Sf9 cells. All transfection reactions were performed using Cellfectin II Reagent. The mock, no DNA treatment (top left panels) and co-transfection of pHRW and pHGW expression plasmids (bottom left panels) were used as controls. Cells were stained with DAPI 72-hours post-transfection, and then visualized using the Cytation 5 Cell Imaging Multi-Mode Reader (BioTek, Winooski, VT) to detect red and green fluorescence. The exposure settings (LED intensity/integration time/camera gain) of the mock were set up as the baseline parameters for image analysis, and treatments were set no more than the mock settings. Cells were visualized with the 40x objective, and each scale bar represents 10 μm.

## Discussion

With the creation of transcriptome sequence resources for *F. occidentalis* and improved proteomics technologies, we have identified the first thrips proteins that bind directly to the TSWV attachment protein, G_N_. With particular relevance to viral attachment to and internalization in epithelia, two TIPs (endocuticle structural glycoprotein, endoCP-G_N_ and cyclophilin) were confirmed to interact directly with G_N_ and were abundant in midgut and salivary gland tissues (21). These data may be the first indication of a protein(s) that serves ‘receptor-like’ roles in transmission biology of the tospoviruses. We narrowed down the G_N_-binding region to the amino terminal region of endoCP-G_N_ excluding the conserved CHB4 domain, setting the stage for future work to decipher the essential amino acids within the non-conserved region necessary to establish the interaction. Three of the TIPs (mATPase, endoCP-V, and enolase) were validated to interact with G_N_ in BiFC assays, but not in MbY2H assays. With regards to other virus activities in host cells, the confirmed affinity of G_N_ with diverse thrips proteins indicates that these insect proteins may be host factors involved in steps in the virus infection cycle in the invertebrate host such as viral replication and/or virion maturation previously observed in both the animal (22, 23) and plant hosts (24, 25). Technical limitations preclude functional analysis of the TIPs in acquisition of virus by larval thrips. Knockdown (RNA interference) and knockout (genome editing) methods have not been developed for larval thrips, even though RNA interference methods have been developed to effectively knockdown genes in the much larger adult female thrips by delivery of dsRNA directly into the hemocoel (26). Using currently available methods, larval thrips do not survive the dsRNA-injection process, and even if successful, knockdown would be delayed thus missing the narrow window of virus acquisition during the early larval development.

The most enriched thrips proteins in the initial screen for those bound to virions or G_N_ (Table 1 and 2, 72%) were cuticular proteins. Cuticular proteins are well characterized as major components of insect hard and soft cuticles (27, 28). Soft cuticles have been documented to line the insect foregut and hindgut (29, 30), and a transmission electron microscopy study documented cuticle lining of the accessory and primary salivary gland (SG) ducts of *F. occidentalis* (31). *In silico* sequence analysis of the three cuticular TIPs (CP-V, endoCP-V, endoCP-G_N_) revealed conserved CHB4 domains (R&R) suggesting their binding affinity to chitin (heteropolymer of N-acetyl-β-D-glucosamine and glucosamine), also a major component of cuticles and peritrophic membranes (PM) lining the midgut epithelium of most insects (32). Hemipteran and thysanopteran midguts lack PMs, and are instead lined with perimicrovillar membranes (PMM) (33, 34) - these structures have been reported to contain lipoproteins, glycoproteins and carbohydrates (35, 36) and more recently, one study documented the occurrence and importance of chitin in the PMM of *Rhodnius prolixus* (kissing bug) midguts, marking the first hemipteran midgut reported to contain chitin (37). Since all three cuticular TIPs were highly expressed in the midgut and SGs of larval *F. occidentalis* in the present study, we hypothesize that chitin or chitin-like structures may impregnate the thrips PMM and SG-linings, forming a matrix with endoCPs - however, this remains to be empirically determined. Alternatively, the thrips TIPs annotated as cuticle proteins with predicted chitin-binding domains may have yet-undescribed functions in insect biology.

Cuticular proteins are emerging as important virus interactors and responders in diverse vector-borne plant virus systems. A CP of the hemipteran vector, *Laodelphax striatellus*, was found to interact with the nucleocapsid protein (pc3) of *Rice stripe virus* (genus *Tenuivirus*, family *Phenuiviridae*) and was hypothesized to be involved in viral transmission and to possibly protect the virus from degradation by a host immune response in the hemolymph (19). Recently, a CP of another hemipteran vector, *Rhopalosiphum padi*, was identified to interact with *Barley yellow dwarf* virus-GPV (genus *Luteovirus*, family *Luteoviridae*) readthrough protein, and the gene transcript of this particular CP was differentially expressed in viruliferous compared to virus-free aphids (38). At the transcript level, thrips cuticular proteins of different types - including the thrips CPRs used in the present study to phylogenetically place the three cuticular TIPs - were identified to be downregulated in TSWV-infected first instar larvae (18). Although the three cuticular TIPs identified in the present study were not reported in the previous study to be differentially-responsive to virus, both implicate cuticle-associated proteins during the early infection events of TSWV in the thrips vector.

Cyclophilins, also known as peptidyl-prolyl cis-trans isomerases, are ubiquitous proteins involved in multiple biological processes, including protein folding and trafficking, cell signaling, and immune responses (39). They have also been shown to promote or prevent virus infection (40, 41), for example, cyclophilin A was found to bind to viral RNA to inhibit replication of *Tomato bushy stunt virus* (genus *Tombusvirus*, family *Tombusviridae*) in plant leaf cells (42), while cyclophilins of the aphid vector *Schizaphis graminum* have been shown to play an important role in *Cereal yellow dwarf virus* (genus *Polerovirus*, family *Luteoviridae*) transmission (43). Interactions between the thrips cyclophilin TIP (with G_N_) documented in the present study may affect similar virus processes, such as virus replication and maturation, or thrips transmission and vector competence (43, 44). The same cyclophilin was determined to be down-regulated in *F. occidentalis* first instar larvae during TSWV infection (45), adding to the body of evidence that viruses modulate expression of cyclophilins (46–51). The cyclophilin interaction with G_N_ was consistent but weak and this may be the reason that it was not observed in the BiFC experiments. Alternative explanations for the discrepancy in the cyclophilin-G_N_ interaction include: the interaction does not occur in plant cells or that the weak interaction was not strong enough to fluoresce over the background to detect an *in planta* interaction. Others have proposed that negative strand virus matrix proteins – structural proteins that package viral RNA - evolved from cyclophilins (52); however, bunyaviruses do not encode a matrix protein. One hypothesis for the direct interaction between the cyclophilin with G_N_ may be to facilitate RNP packing into the virus particle, perhaps serving as a surrogate matrix protein for TSWV.

Like cyclophilins, enolases of diverse hosts have been identified as both responsive to and interactive partners with viruses. In general, enolases are essential metalloenzymes that catalyze the conversion of 2-phosphoglycerate (2-PGE) to phosphoenolpyruvate (PEP) in the glycolytic pathway for energy metabolism (53). Some are matrix metalloproteases known to cleave cell surface receptors, modulate cytokine or chemokine activities, or release apoptotic ligands by degrading all types of extracellular matrix proteins, such as collagen, elastin, fibronectin, laminin, gelatin, and fibrin (54). The enolase TIP identified in the present study was previously reported to be up-regulated in L1 bodies infected with TSWV (45), as was the case for enolase in response to RSV in bodies of the planthopper vector, *L. striatellus* (55). In the case of flaviviruses, *Aedes aegypti* enolase was shown to directly interact with purified virus and recombinant envelope GP of dengue virus (56) and West Nile virus envelope protein (57). The localization of this enolase in brush border membrane vesicles of this mosquito species (58) strengthens the case for a proposed receptor role in virus entry into vector mosquito midguts. Other insect-virus studies have proposed a role for enolase in antiviral defense (54) and tracheal basal laminal remodeling aiding in virus escape from the gut (59). If remodeling of the midgut basal lamina via enolase interactions occurs in TSWV-infected larval thrips, that could be one hypothesis supporting dissemination of TSWV from the larval midgut into the principal SGs (15).

The other TIP known to play a role in energy production is mitochondrial ATP synthase α subunit. The multi-subunit enzyme mATPase is responsible for generating the majority of cellular ATP required by eukaryotes to meet their energy needs. As with the other non-cuticle TIPs, mATPase α subunit was previously identified to be differentially-abundant (up-regulated) under TSWV infection (45), as was the case for RSV-infected *L. striatellus* vector planthoppers (55). Mitochondria have also been previously implicated in virus-host biology. For example, *African swine fever virus* (genus *Asfivirus*, family *Asfarviridae*) has been shown to induce migration of mitochondria to the periphery of viral factories (60), possibly suggesting that mitochondria supply energy for viral morphogenetic processes. The finding that two TIPs in the present study have ontologies in energy production and metabolism suggests that perturbation or direct interactions with these host proteins may be required for the successful infection of *F. occidentalis* by TSWV.

The discovery of six TIPs is a significant step forward for understanding thrips interactions with tospoviruses. The first evidence of TSWV protein-thrips protein interactions was presented 20 years ago (61) and the proteins described herein are the first thrips proteins documented to interact directly with the viral glycoprotein, G_N_, involved in virus attachment to the midgut epithelial cells of the insect vector. In other eukaryotes, the six interacting proteins have biological functions that point to their putative roles in facilitating the virus infection/replication cycle by acting as a receptor or other essential step in the virus life cycle and/or host-response via a defense mechanism. The virus-host systems that have defined functions for analogous TIPs include plant viruses, arboviruses, and animal/human viruses, and the findings described here provide a framework for further exploration and testing of new hypotheses regarding their roles in TSWV-thrips interactions.

## Materials and methods

### Insect rearing and plant and virus maintenance

The *F. occidentalis* colony was established from insects collected on the island of Oahu, HI, and was maintained on green beans (*Phaseolus vulgaris*) at 22°C (± 2°C) under laboratory conditions as previously described (62). Thrips were age-synchronized based on their developmental stages. For the localization and bimolecular fluorescence complementation (BiFC) experiments, wildtype and transgenic *Nicotiana benthamiana* expressing CFP:H2B or RFP:ER (63) were grown in a growth chamber at 25°C with a 14-hour light at 300 μM intensity and 10-hour dark cycle. TSWV (isolate TSWV-MT2) was maintained by both mechanical inoculation and thrips transmission using *Datura stramonium* and *Emilia sonchifolia*, respectively (12). To avoid generation of a virus isolate with an insect transmission deficiency, the virus was mechanically passaged only once. The single-pass mechanically-inoculated symptomatic *D. stramonium* leaves were used for insect acquisition of TSWV. Briefly, synchronized *F. occidentalis* first instar larvae (0-17-hour old) were collected and allowed an acquisition access period (AAP) on *D. stramonium* for 24 hours. After acquisition, *D. stramonium* leaves were removed and these larvae were maintained on green beans until they developed to adults. Viruliferous adults were transferred onto clean *E. sonchifolia* for two days. After inoculation, thrips and inoculated *E. sonchifolia* plants were treated with commercial pest strips for two hours before the plants were moved to the greenhouse for TSWV symptom development. The thrips-transmitted, TSWV symptomatic *E. sonchifolia* leaves were only used for mechanical inoculation.

### TSWV purification

Mechanically inoculated *D. stramonium* leaves were used for TSWV purification via differential centrifugation and a sucrose gradient. Symptomatic leaves were homogenized in extraction buffer (0.033 M KH_2_PO_4_, 0.067 M K_2_HPO_4_, and 0.01 M Na_2_SO_3_) in a 1:3 ratio of leaf tissue to buffer. The homogenate was then filtered through four layers of cheesecloth, and the flow through was centrifuged at 7,000 rpm (7,445 *g*) for 15 min using the Sorvall SLA 1500 rotor. To remove the cell debris, the pellet was resuspended in 65 mL 0.01 M N_2_SO_3_ and was centrifuged again at 8,500 rpm (8,643 g) for 20 min using the Sorvall SS34 rotor. The supernatant that contained the virions was centrifuged for 33 min at 29,300 rpm (88205 *g*) using the 70 Ti rotor, and the pellet was resuspended in 15 mL 0.01 M Na_2_SO_3_ followed by another centrifugation at 9,000 rpm (9,690 *g*) for 15 min using the Sorvall SS34 rotor. The centrifugation series was repeated one additional time. The pellet was resuspended and loaded on a sucrose gradient (10 to 40% sucrose), which was centrifuged for 35 min at 21,000 rpm (79,379 *g*) using the SW28 rotor. The virion band was collected and centrifuged for 1 hour at 29,300 rpm (88,205 *g*) using the 70 Ti rotor. The pellet was resuspended in 100 to 200 μl of 0.01 M Na_2_SO_3_. All centrifugation steps were performed at 4°C to prevent virion degradation. The purified virus was quantified using the bicinchoninic acid (BCA) protein assay kit (ThermoFisher Scientific, Waltham, MA, USA) following the manufacturer’s instructions.

### *F. occidentalis* L1 total protein extraction, quantification and two-dimensional (2-D) electrophoresis

Total proteins from age-synchronized healthy larval thrips (0-17-hour old) were extracted using the trichloroacetic acid-acetone (TCA-A) method (64, 65). Briefly, whole insects were ground using liquid nitrogen, and were dissolved in 500 μl TCA-A extraction buffer (10% of TCA in acetone containing 2% β-mercaptoethanol). This mixture was incubated at −20°C overnight and centrifuged at 5,000 *g*, 4°C for 30 min. After 3 washes with ice-cold acetone then air-drying, the pellet was resuspended in 200 μl General-Purpose Rehydration/Sample buffer (Bio-Rad Laboratories, Hercules, CA, USA). The suspension was centrifuged at 12,000 *g* for 5 min and the protein supernatant was quantified using the BCA protein assay kit (ThermoFisher Scientific) following manufacturer’s instructions. For each gel, 150 μg of total protein extract was applied to an 11-cm IPG strip (pH 3–10) for isoelectric focusing (IEF). The IEF, IPG strip equilibration and second dimension separation of proteins were performed under the same conditions described by Badillo-Vargas et al (45).

### Overlay assays

To identify thrips proteins that bind to TSWV virions and recombinant glycoprotein G_N_, we conducted gel overlay assays. For the purified virion overlays, the experiment was performed four times (biological replications); and for the G_N_ overlay, the experiment was performed twice. For probing the protein-protein interactions, each unstained 2-D gel was electro-transferred onto Hybond-C Extra nitrocellulose membrane (Amersham Biosciences, Little Chalfont, UK) overnight at 30 V (4°C) in protein transfer buffer (48 mM Tris, 39 mM glycine, 20% methanol, and 0.037% SDS). Then, the membrane was incubated with blocking buffer (PBST containing 0.05% Tween 20 and 5% dry milk) for 1h at room temperature on a rocker with a gentle rotating motion. Three different antigens were used to probe the thrips protein membranes: purified TSWV virions, recombinant glycoprotein G_N_ (*E. coli* expressed), and virus-free plant extract from a mock virus purification (negative control). An additional negative control blot (no overlay) treated with antibodies alone was included in each overlay replicate. For the virus and G_N_ treatments, 25 μg/mL and 3.5 μg/mL of purified TSWV virions and recombinant G_N_ glycoprotein, respectively, were incubated with membranes in blocking buffer at 4°C overnight with gentle rotating motion. Membranes were washed three times using PBST and were incubated with polyclonal rabbit anti-TSWV G_N_ antiserum at 1:2,000 dilution in blocking buffer for 2 hours at room temperature (9, 21). After washing with PBST, membranes were incubated with HRP-conjugated goat-anti rabbit antiserum at 1:5,000 dilution in blocking buffer for 1 hour at room temperature. The ECL detection system (Amersham Biosciences) was used for protein visualization following the manufacturer’s instructions. The protein spots that were consistently observed on the membranes were first compared with those proteins spots that interacted with antibody-only blots (Fig 1A and Fig 2A) and virus-free plant extract blots, and then they were pinpointed on the corresponding Coomassie Brilliant Blue G-250-stained 2-D gels for spot picking.

### Identification of TIPs

Protein spots that were consistently identified in the 2-D gel overlays were selected and manually picked for analysis. The picked proteins were processed and subjected to ESI mass spectrometry as previously described (21). Protein spots (peptides) that had Mascot scores (Mascot v2.2) with significant matches (*P* ≤ 0.05) to translated *de novo*-assembled contigs (all six frames) derived from mixed stages of *F. occidentalis* (“Fo Seq” 454-Sanger hybrid) (45) were identified and NCBI Blastx was performed on the contigs to provisionally annotate (*E* < 10^−10^) the protein and to predict conserved motifs using the contig as the query and the NCBI non-redundant protein database as the subject.

A second round of TIP candidate selection was conducted for stringency in moving forward to cloning and confirmation of interactions. A contig sequence was retained if it contained a complete predicted ORF (*i. e*., presence of both start and stop codons predicted with Expasy, Translate Tool, http://web.expasy.org/translate/) and had at least 10% coverage by a matching peptide(s) identified for a spot as predicted by Mascot. *i. e*., removal of proteins identified by a single peptide with less than 10% coverage to a *Fo* Seq contig and/or contigs with incomplete ORFs (lacking predicted stop codon). The translated ORFs were queried against the NCBI non-redundant protein database (Blastp), and CCTOP software (http://cctop.enzim.ttk.mta.hu) (66) and SignalP 4.1 Server (http://www.cbs.dtu.dk/services/SignalP/) (67) were used to predict the presence of transmembrane domains and signal peptides, respectively. Prosite (http://prosite.expasy.org/) was used to analyze putative post-translational modifications that may have affected electrophoretic mobility of identical proteins in the overlay assays, *i. e*., same peptide sequence or *Fo* Seq contig match identified for more than one protein spot.

### Classification and phylogenetic analysis of the three confirmed cuticular TIPs

Given the apparent enrichment of putative cuticular proteins (CP) identified in the overlay assays and the subsequent confirmation of three of those TIPs (CP-V, endoCP-G_N_, endoCP-V), it was of interest to perform a second layer of protein annotations. The ORFs (amino acid sequence) of the three confirmed CP TIPs, 19 exemplar insect orthologous sequences obtained from NCBI GenBank, and a significant collection of structural CP transcripts previously reported to be differentially-expressed in TSWV-infected larval thrips of *F. occidentalis* (18) were subjected to two complementary arthropod CP prediction tools. CutProtFam-Pred (http://aias.biol.uoa.gr/CutProtFam-Pred/home.php) (68) was used to classify each amino acid sequence by CP family – there are 12 described families for arthropods, each distinguished by conserved sequence motifs shared by members (28) – and CuticleDB (http://bioinformatics.biol.uoa.gr/cuticleDB) (69) was used to distinguish what was found to be two enriched, chitin-binding CP families in our dataset: CPR-RR1 and CPR-RR2, *i. e*., R&R Consensus motif (17). The sequences flanking the RR1 and RR2 predicted chitin-binding domains were so divergent between the thrips CPs and across the entire set of CPs (thrips and other insects) that alignments using full-length ORFs were ambiguous and uninformative, thus illustrating the utility of the R&R Consensus for inferring evolutionary history of CP proteins. The flanking sequences were trimmed manually, and the R&R consensus sequences (RR1 and RR2) were aligned with MEGA7 (70) using ClustalW. Phylogenetic analyses were performed in MEGA7 using the Neighbor-Joining (NJ) method and the best substitution models determined for the data - Dayhoff matrix-based (71) or Jones Taylor Thorton (JTT) (72) methods for amino acid substitutions with Gamma distribution - to model the variation among sites. Bootstrap consensus trees (500 replicates) were generated by the NJ algorithm with pairwise deletion for handling gaps. The analysis involved 46 sequences and there were 95 amino acid positions in the final dataset.

### Cloning of candidate TIPs and TSWV genes

For generation of full-length clones of TIPs that were used in various protein-protein assays, total RNA was extracted from L1 thrips (0-17-hour old) using 1 mL Trizol Reagent (ThermoFisher Scientific), then 200 μl chloroform and was precipitated with 500 μl isopropanol. The RNA pellet was dissolved in nuclease-free water, and 1μg total RNA was used for cDNA synthesis using the Verso cDNA Synthesis kit (ThermoFisher Scientific). The PCR was performed to amplify six identified TIP ORFs using high fidelity polymerase, FailSafe (Epicentre, Madison, WI, USA). The designed primers used are listed in Table S2. Amplicons were cloned into pENTR-D/TOPO (ThermoFisher Scientific).

TSWV genes were also cloned to pENTR-D/TOPO, then recombined to different vectors using Gateway cloning techniques. Coding sequences of different glycoprotein forms (soluble (G_N_-S) and insoluble (G_N_)) were amplified from pGF7 (73). Primers used for PCR were listed in Table S2.

### Polyclonal antisera against TIPs

To generate antibodies to the TIPs, the protein sequence was analyzed for multiple features such as antigenicity and hydrophobicity by the antibody manufacturer (GenScript, Piscataway, NJ), using the OptimumAntigen^™^ Design Tool (https://www.genscript.com/antigen-design.html). For each TIP, a 14 amino acid peptide was selected based on these predictions and by sequence alignments to other predicted protein sequences in GenBank. Due to the conserved CHB4 domain in endoCP-G_N_, endoCP-V, and CP-V, the polyclonal antibodies against these three TIPs were generated using their non-conserved region. The peptides were synthesized, and all antisera were produced using mice (GenScript, Piscataway, NJ, United States). The peptide sequences for each TIP that were used for the antibody generation were: cyclophilin, LESFGSHDGKTSKK; enolase, ELRDNDKSQYHGKS; CP-V, TDSGQYRKEKRLED; endoCP-G_N_, STKVNPQSFSRSSV; endoCP-V, VNPDGSFQYSYQTG; and mATPase, GHLDKLDPAKITDF.

### Validation of antisera specificity against each TIP (peptide) using dot blot

Peptide antibodies to the six TIPs were generated by GenScript using their standardized work flow. For each TIP, the amino acids of highest antigenic potential were identified, peptides were synthesized, antibodies were generated by injection in mice, and antibodies were tested for reactivity and specificity using dot-blot assays. The peptides that were used to generate each antibody were diluted to 100 μg/mL using 1×PBS (pH=7.2), with the exception of the CP-V peptide that was diluted to 2.5 mg/mL (briefly explain why – lower sensitivity?). Two ul of each diluted peptide was spotted onto the same nitrocellulose membrane strip along with the controls of PBS and pre-immune serum (500,000 × dilution). A total of 6 membrane strips, one for each TIP-peptide antibody, were loaded with the same peptide samples and controls. After membrane strips dried, each strip was incubated with blocking buffer (5% non-fat milk in TBS-T), followed by incubation with the six different primary antibodies (0.5 μg/mL, produced by GenScript), respectively. After three washes with TBS-T (3×10 min), all membrane strips were incubated with secondary antibody, goat anti-mouse IgG (H+L)-HRP conjugate (1:5,000 dilution, Bio-Rad Laboratories). After three washes with TBS-T, the SuperSignal™ West Dura Extended Duration Substrate (ThermoFisher Scientific) was added onto individual membrane strips. Each membrane strip was developed separately for 5 to 10 min, however, the membrane strip that was incubated with the CP-V peptide antibody was developed for 40 min. Then a picture was taken using iBright Imaging system (CL1000, ThermoFisher Scientific). The blocking, primary and secondary antibody incubation steps were incubated for 1 hour at room temperature, and the strip probed with CP-V peptide antibody was incubated for 2 hours at room temperature. The entire experiment was performed three times.

### Immunolabeling thrips guts, Malphigian tubules, and salivary glands

To determine the location of TIPs expression in the most efficient thrips stage that acquires TSWV (L1), we used the TIPs antibodies in immunolocalization experiments. Treatments in the experiments included peptide antibodies to the TIPs and background controls of dissected insects incubated with i) only secondary antibody and ii) insects treated with pre-immune serum and secondary antibody. Newly emerged larvae (0-17-hour old) were collected from green beans and were then fed on 7% sucrose solution for 3 hours to clean their guts from plant tissues. The larvae were dissected on glass slides using cold phosphate saline (PBS) buffer and Teflon coated razor blades. The dissected thrips were transferred into 2-cm-diam., flat-bottomed watch glasses (U.S. Bureau of Plant Industry, BPI dishes) and the tissues were fixed for 2 hours using 4% paraformaldehyde solution in 50 mM sodium phosphate buffer (pH 7.0). The tissues were washed using PBS buffer after fixation and were incubated with PBS buffer including 1% Triton X-100 overnight. The overnight permeabilized tissues were then washed before incubation in blocking buffer which included PBS, 0.1% Triton X-100 and 10% normal goat serum (NGS) for 1 hour. After removing the blocking buffer, the dissected thrips were incubated with primary antibody, 100 μg/mL mice-generated antisera against each individual TIP (GenScript) that was diluted in antibody buffer (0.1% Triton X-100 and 1% NGS). After washing, 10 μg/mL secondary antibody, goat anti-mouse antibody conjugated with Alexa Fluor 488 (ThermoFisher Scientific) was used to incubate the dissected thrips organs. Incubation was performed at room temperature for 2.5 hours, 1x PBS buffer was used for washing and every wash step included three rinses, and the secondary antibody incubation was protected from light by covering the samples with aluminum foil. After removing antibodies and washing, dissected thrips were incubated for 2 hours with Phalloidin-Alexa 594 conjugated (ThermoFisher Scientific) in 1x PBS with a concentration of 4 units/mL for actin staining. After washing, the tissues were transferred onto glass slides, and SlowFade™ Diamond Antifade Mountant with DAPI (ThermoFisher Scientific) was added onto tissues to stain the nuclei. The cover slips were slowly placed on tissues to avoid bubbles, then sealed with transparent nail polish at the edges. After blocking, the dissected thrips tissues that were only incubated with secondary antibody (without adding primary antibody) and the tissues incubated with each pre-immune mouse antiserum (GenScript) were used as negative controls, respectively. All the experiments were performed twice.

Inherent with very small tissues (< 1 mm body size), there were common losses or damaged tissues during the dissection process and staining procedures; so only the number of visibly intact tissue that made it through to microscopic observation were used for data collection and this number varied for each type of tissue (Table S4). The auto-fluorescent background from thrips tissues incubated with each pre-immune antiserum and secondary antibodies was slightly higher than the thrips tissues incubated with PBS buffer and secondary antibodies (negative control) (data not shown), therefore, the confocal laser settings (power and percent gain) were adjusted to remove any background fluorescence observed for these treatments.

### Split-ubiquitin membrane-based yeast two-hybrid (MbY2H)

The MbY2H system was used to validate TSWV G_N_-TIPs interactions identified in the gel overlay assays. The MbY2H system enables validation of interactions for soluble and integral membrane proteins. TSWV G_N_ coding sequence were cloned into the MbYTH vector pBT3-SUC, and the six TIP ORFs were cloned to vector pPR3N using the SfiI restriction site (Dualsystems Biotech, Schlieren, Switzerland). To identify the region of endoCP-G_N_ that binds to TSWV G_N_ using MbY2H, the amino acid sequence of endoCP-G_N_ (284aa) was used to search against the NCBI non-redundant protein database using Blastp. The conserved CHB4 domain was located at the C-terminus of endoCP-G_N_ (amino acid 190-246). Therefore, the possible interacting domains, the non-conserved region of endoCP-G_N_ (1-189aa) and the conserved CHB4 domain (190-274aa), were individually cloned into pPR3N using the SfiI restriction site. Based on the Blastp results, the homologous sequences from other insect species encompassed some additional amino acids upstream of the CHB4 domain; therefore, we made an alternative construct that included the conserved CBH4 domain starting from amino acid 177. Hence, the coding sequence of 1-176aa and 177-284aa of endoCP-G_N_ were also cloned to pPR3N using the SfiI restriction site. Primers used for cloning are listed in Table S3.

The MbY2H assays were performed using the manufacturer’s instructions with recombinant plasmids that were confirmed by Sanger sequencing. Yeast (strain NYM51) competent cells were freshly prepared and recombinant bait plasmids, pBT3-SUC-G_N_ were transformed into yeast cells. Briefly, 1.5 μg of bait plasmids were added into 100 μl of yeast competent cells with 50 μg of denatured Yeastmaker Carrier DNA (Takara Bio USA, Mountain View, CA) and 500 μl PEG/LiAc. The mixture was incubated at 30°C for 30 min with mixing every 10 min. Twenty μl of DMSO was then added into each reaction, and the cells were incubated at 42°C for 20 min with mixing every 5 min. After centrifugation at 14,000 rpm for 15 sec, the supernatant was removed, and the pellet was resuspended in 1 mL of YPDA media. The re-suspended cells were incubated at 30°C for 90 min with shaking at 200 rpm. Then, cells were centrifuged at 14,000 rpm for 15 sec, and resuspended in 500 μl of sterile 0.9% (w/v) NaCl, which was then spread and cultured on SD/–Trp dropout media at 30°C until the colonies were visible. Several colonies from the same SD/–Trp plate were cultured for preparing yeast competent cells. Then each individual recombinant plasmid, pPR3N-TIP or pPR3N-partial endoCP-G_N_ (1.5 μg/transformation reaction), was transformed into yeast competent cells expressing fused Nub-G_N_. The transformants were cultured on both SD/–Leu/–Trp double dropout (DDO) and SD/–Ade/–His/–Leu/–Trp quadruple dropout (QDO) media. The positive controls included transformation of pOst1-NubI into the yeast strain NYM51 that already expressed fused Nub-G_N_ or Nub-N, as well as co-transformation of pTSU2-APP and pNubG-Fe65 into the yeast strain NYM51. Transformation of pPR3N (empty vector) into the yeast strain NYM51 that already expressed fused Nub-G_N_ was used as the negative control. Interactions between G_N_-Cub and NubI, G_N_-Cub and NubG were used as positive and negative controls respectively. All transformants were spread and cultured on both DDO and QDO media and cultured at 30°C in an incubator. The entire experiment was performed three times.

### Yeast β-galactosidase assay

Expression of the reporter gene *LacZ* and the activity of expressed β-galactosidase in yeast cells derived from MbY2H was determined by a β-galactosidase assay kit following the manufacturer’s protocol (ThermoFisher Scientific). Each yeast colony was transferred, mixed with 250 μl of Y-PER by vortex, and their initial OD_660_ value was determined. After adding 250 μl 2X β-galactosidase assay buffer to the mixed solution, the reaction was incubated at 37°C until the color change of solution was observed. Two hundred μl of β-galactosidase assay stop solution was added immediately into color change solution, and the reaction time was recorded. Cell debris was removed by centrifugation at 13,000 *g* for 30 seconds. Supernatant was transferred into cuvettes to measure OD_420_ using the blank including 250 μl of Y-PER reagent, 250 μl β-galactosidase assay buffer and 200 μl β-galactosidase assay stop solution. The β-galactosidase activity was calculated using the equation from the manufacturer’s protocol.

### GFP fusion protein expression and bimolecular fluorescence complementation (BiFC) in *Nicotiana benthamiana*

To visualize protein expression and localization in plants, TSWV G_N_ (ORFs G_N_ and G_N_S) and TIP ORFs (mATPase, CP-V, endoCP-V, endoCP-G_N_, cyclophilin and enolase) were expressed as fusions to autofluorescent proteins. They were moved from their entry clones into pSITE-2NB (GFP fused to the carboxy terminus of the protein of interest) or pSITE-2CA (GFP fused to the amino terminus of the protein of interest) using Gateway LR Clonase (74). After validation of plasmids by Sanger sequencing, they were transformed into *Agrobacterium tumefaciens* strain LBA 4404. The transformed LBA 4404 was grown for two days at 28°C and re-suspended in 0.1 M MES and 0.1 M MgCl_2_ to an OD_600_ between 0.6 to 1. After the addition of 0.1 M acetosyringone, the suspension was incubated at room temperature for two hours, and then infiltrated in transgenic *N. benthamiana* expressing an endoplasmic reticulum (ER) marker fused to the red fluorescent protein (m5RFP-HDEL) (63). Two days after infiltration, leaf tissue was mounted in water on a microscope slide for detection of GFP by confocal microscopy. Plants were infiltrated a minimum of two separate occasions with at least two leaves per plant in two different plants. A minimum of fifty cells were visualized in each plant to confirm the localization patterns of the proteins *in planta*.

The preliminary localization results and sequence analysis informed the fusion construct design for BiFC assays. Signal peptides were identified in the amino terminus of G_N_, and three TIPs (all cuticle proteins) and the signal peptide is required for proper localization and function of fusion-GFP/YFP proteins in *N. benthamiana* for BiFC assays. Based on the expression and localization results of GFP fusion proteins, we fused half YFPs (either amino or carboxy half of YFP) to the carboxy termini of all proteins with N-terminal signal peptides using BiFC plasmids pSITE-NEN and pSITE-CEN (63). All ORFS were transferred between plasmids using Gateway LR Clonase II Enzyme Mix (ThermoFisher Scientific). All clones were transformed into *A. tumefaciens* strain LBA 4404 and confirmed by Sanger sequencing.

Each combination of TIPs and TSWV G_N_ and G_N_-S was infiltrated into *N. benthamiana* expressing CFP fused to a nuclear marker, histone 2B, (CFP-H2B) (63), and a minimum of three independent experiments with two plants and two leaves per plant for each combination of proteins. For the analysis of interactions, a minimum of 50 cells with similar localization patterns was required to confirm the interaction and a minimum of two separate images were captured on each occasion for documentation. GST fusions to YFP halves were utilized as a non-binding control for each of the TIPs. To be recorded as a positive interaction, fluorescence of the interacting TSWV protein-TIPs was required to be above that observed between each TIP and GST.

### Laser scanning confocal microscopy

Confocal microscopy was used to detect the fluorescent signal produced from TIP antibody labelling in thrips tissues and BiFC experiments in plants. All images were acquired on a Zeiss LSM 780 laser scanning confocal microscope using the C-Apochromat 40x/1.2 W Korr M27 and Plan-Apochromat 20x/0.8 M27 objectives. Image acquisition was conducted on Zen 2 black edition v. 10.0.0 at 1024 × 1024 pixels with a scan rate of 1.58 μs per pixel with pixel average of 4-bit and 16-bit depth. The laser power and percent gain settings for detection of nuclei and actin as well as the bright field were adjusted accordingly. Laser power and percent gain settings for detection of TIPs were equal or smaller than their controls. Z-stacks were taken for localization of TIPs in thrips. Eight (TIPs localization) Z-stack slides were processed using Maximum intensity projection using Zen 2 black. Zen 2 blue edition lite 2010 v. 2.0.0.0 was used for image conversion to jpeg format.

### Co-localization of TIPs and TSWV G_N_ in insect cells

The ORFs of cyclophilin, endoCP-G_N_ and TSWV G_N_ were cloned into pENTR/D-TOPO (Thermo Fisher Scientific, Grand Island, NY). TSWV G_N_ was amplified by primers ENTR-endoCP-G_N_F and ENTR-TSWV-G_N_R1353 (Table S2). The cyclophilin and endoCP-G_N_ ENTR clones were recombined into pHWR, and the TSWV G_N_ ENTR clone was moved into pHWG (*Drosophila* gateway collection, DGRC, Bloomington, Indiana). Both pHWR and pHWG have Hsp70 promotor and gateway cloning cassette, and both RFP and GFP were expressed at the C-terminus of TIPs or TSWV G_N_.

The recombinant expression constructs were confirmed by Sanger sequencing and then transfected into Sf9 cells. Single- or co-transfections were performed using Cellfectin II Reagent (ThermoFisher Scientific) following the manufacturer’s protocol. Briefly, Sf9 cells were counted, diluted to 5×10^5^, and then 2 ml aliquots were seeded into each well of a 6-well plate. Eight μL of Cellfectin II reagent and 3 ng of each recombinant plasmid were diluted in 100 μL Grace’s medium (Thermofisher Scientific), respectively. After vortex-mixing, both diluted DNA and diluted Cellfectin II reagent were incubated at room temperature for 30 min and then were combined and incubated for an additional 30 min. Another 800 μL of Grace’s medium was added into each DNA-lipid mixture, and the entire 1 mL solution was slowly added onto Sf9 cells. The transfection mix was incubated for 5 hours at 27°C after which the solution was removed and replaced by 2 mL Sf-900 III medium. Single transfected plasmids were pHWR-cyclophilin, pHWR-endoCP-G_N_ and pHWG-TSWV G_N_; co-transfected plasmids were pHWR-cyclophilin and pHWG-TSWV G_N_; pHWR-endoCP-G_N_ and pHWG-TSWV G_N_. To rule out non-specific interactions between the proteins of interests (TIPs or G_N_) and the autofluorescent protein tags, they were co-transfected with unfused RFP or GFP. In addition, a mock (no DNA) transfection was also included as negative control.

After 72 h, Sf9 cells were resuspended and re-seeded in a 24-well glass bottom Sensoplate (Greiner Bio-One, Monroe, NC) with 1:2 dilution. The cells were stained with DAPI and then visualized by the Cytation 5 Cell Imaging Multi-Mode Reader with objectives 40x PL FL and 20x PL FL (BioTek, Winooski, VT). Image acquisition was performed with BioTek Gen 5 Microplate Reader and Imager Software, version 3.04. Images were captured using default settings. To detect GFP and RFP, the exposure settings (LED intensity/integration time/camera gain) of mock transfected cells were set up as the baseline and different treatments were set to no more than the mock settings. Other parameter settings for detection of nuclei and bright field were adjusted accordingly. The entire experiment was performed four times.

## Data availability

The GenBank accession numbers for six TIPs are: cyclophilin, MH884760; enolase, MH884759; cuticular protein: CP-V, MH884758; endoCP-G_N_, MH884757; endoCP-V, MH884756; mitochondrial ATP synthase α, MH884761.

## Acknowledgements

We thank Thomas L. German and Ranjit Dasgupta for providing purified G_N_-S for protein overlays. This project was supported by the following grants: USDA-NIFA 2007-35319-18326 and 2016-67013-27492, USDA-FNRI 6034-22000-039-06S, and National Science Foundation CAREER Grant IOS-0953786. Ismael E. Badillo-Vargas was partially supported by the National Institute of Food and Agriculture Predoctoral Fellowship, grant KS602489.

## Supporting Information

**Fig. S1 Phylogenetic analysis of cuticle protein (CP) R&R consensus sequences in first instar larvae of *F. occidentalis*.** The Neighbor-Joining (NJ) method was performed with the Jones Taylor Thorton (JTT) matrix-based method for amino acid substitutions with Gamma distribution to model the variation among sites. The bootstrap consensus tree (500 replicates) was generated by the NJ algorithm with pairwise deletion for handling gaps. Branches corresponding to partitions reproduced in less than 70% bootstrap replicates were collapsed. The numbers shown next to branches indicate the percentage of replicate trees in which the associated taxa (sequences) clustered together in the bootstrap test. The analysis involved 46 sequences – the three cuticular TSWV-interacting proteins (TIPs: cuticle protein-V (CP-V), endocuticle structural glycoprotein-V (endoCP-V) and endocuticle structural glycoprotein-G_N_ (endoCP-G_N_) (blue text), the ‘gold-standard’ Pfam database extended R&R consensus sequence (pf00379), 19 insect orthologous sequences obtained from NCBI GenBank, and 23 structural CPs and endoCPs (translated transcripts, designated with FOCC or CUFF identifiers) previously reported to be differentially-expressed in whole bodies of TSWV-infected L1s of *F. occidentalis* (27). There were 95 amino-acid positions in the final dataset. RR1 and RR2 = <u>C</u>uticle <u>P</u>rotein <u>R</u>ebers and <u>R</u>iddiford (CPR family, RR1 and RR2 types) extended consensus, a conserved chitin-binding motif (chitin_bind_4 = CHB4).

**Fig. S2 Individual channels of immune-labeled TSWV-interacting proteins (TIPs) within first instar larvae of *F. occidentalis*.** The synchronized first instar larvae (0-17-hour old) were dissected and immunolabeled using specific antibodies against each TIP as indicated. Thrips tissues incubated with pre-immune mouse antiserum as controls are depicted here. Confocal microscopy detected green fluorescence (Alexa Fluor 488) that represents the localization of each TIP, red represents Alexa Fluor 594 labeled actin; blue represents DAPI labeled nuclei. All scale bars are equal to 50 μM.

**Fig. S3 Localization of TSWV-interacting proteins (TIPs) fused to green fluorescent protein (GFP) in *Nicotiana benthamiana*. Plants** transgenic for an RFP-ER marker were infiltrated with *Agrobacterium tumefaciens* strain LBA 4404 suspensions of TIPs constructs. Each row indicates the specific TIP-GFP fusion in relation to the RFP-ER marker. The columns are as follows: GFP channel, RFP channel and the overlay between the two channels. All scale bars are equal to 20 μm.

**Fig. S4 Co-expression of fusion proteins in Sf9 insect cells.** Three control sets of recombinant plasmids were co-transfected into insect Sf9 cells to ensure that unfused autofluorescent proteins do not alter protein localization (cyclophilin, endo-CP-G_N_, and G_N_), pHWR-cyclophilin and pHGW (expression of cyclophilin-RFP and GFP); pHWR-endoCP-G_N_ and pHGW (expression of endoCP-G_N_-RFP and GFP); pHRW and pHWG-TSWV G_N_ (expression of RFP and TSWV G_N_-GFP). All transfection reactions were performed using Cellfectin II Reagent, and cells were stained with DAPI at 72 hours post transfection. The red and green fluorescence were detected by Cytation 5 Cell Imaging Multi-Mode Reader. The exposure settings (LED intensity/integration time/camera gain) of mock were set up as the baseline, and different treatments were set no more than the mock settings. Each scale bar represents 10 μm.

