## Supplemental tables 1-4 for "Discovery of novel thrips vector proteins that bind to the viral attachment protein of the plant bunyavirus, tomato spotted wilt virus"

Table S1, Confirmation of interactions between TSWV G_N_ and two TIPs, cyclophilin and endoCP-G_N_, as well as different regions of endoCP-G_N_ using β-galactosidase assay. The volume (V) of each reaction was 0.25ml, and the β-galactosidase activity was calculated using the equation following the manufacturer’s protocol. β-galactosidase activity = (1000*A_420_)/(time*V*OD_660_).

| Protein-protein | Replicate | OD_660_ | A_420_ | Time of incubation (in minutes) | Activity |
| --- | --- | --- | --- | --- | --- |
| G_N_-endoCP-G_N_ | 1 | 0.22 | 0.328 | 3.25 | 1987.879 |
|  | 2 | 0.079 | 0.335 | 1.75 | 11308.02 |
|  | 3 | 0.031 | 0.21 | 1 | 36129.03 |
|  | 4 | 0.062 | 0.414 | 2.75 | 10683.87 |
|  | 5 | 0.067 | 0.536 | 2.75 | 12800 |
|  | 6 | 0.046 | 0.103 | 6.25 | 1492.754 |
|  | 7 | 0.084 | 0.534 | 1.25 | 25428.57 |
|  | 8 | 0.059 | 0.308 | 4.25 | 5220.339 |
|  | 9 | 0.063 | 0.501 | 2.75 | 12723.81 |
| G_N_-cyclophilin | 1 | 0.057 | 0.212 | 2.25 | 7438.596 |
|  | 2 | 0.116 | 0.754 | 1 | 34666.67 |
|  | 3 | 0.111 | 0.534 | 1.75 | 12828.83 |
|  | 4 | 0.032 | 0.304 | 2.25 | 16888.89 |
|  | 5 | 0.069 | 0.504 | 0.75 | 38956.52 |
|  | 6 | 0.24 | 0.494 | 0.75 | 10977.78 |
|  | 7 | 0.064 | 0.339 | 2.25 | 9416.667 |
| G_N_-endoCP-G_N_1-176 | 1 | 0.049 | 0.075 | 4.08 | 1500.60 |
|  | 2 | 0.029 | 0.076 | 6.67 | 1571.63 |
|  | 3 | 0.037 | 0.081 | 6.08 | 1440.26 |
|  | 4 | 0.041 | 0.373 | 0.95 | 38305.52 |
|  | 5 | 0.029 | 0.311 | 2 | 21448.28 |
|  | 6 | 0.037 | 0.27 | 1.18 | 24736.60 |
|  | 7 | 0.027 | 0.105 | 6.5 | 2393.16 |
|  | 8 | 0.023 | 0.073 | 5.25 | 2418.22 |
|  | 9 | 0.036 | 0.071 | 3.75 | 2103.70 |
| G_N_-endoCP-G_N_1-189 | 1 | 0.045 | 0.124 | 2.25 | 4898.77 |
|  | 2 | 0.041 | 0.135 | 2.75 | 4789.36 |
|  | 3 | 0.022 | 0.112 | 3.33 | 6115.21 |
|  | 4 | 0.042 | 0.263 | 0.83 | 30177.85 |
|  | 5 | 0.012 | 0.292 | 1.92 | 50694.44 |
|  | 6 | 0.036 | 0.219 | 1.08 | 22530.86 |
|  | 7 | 0.044 | 0.161 | 3 | 4878.79 |
|  | 8 | 0.039 | 0.133 | 5.78 | 2360.04 |
|  | 9 | 0.046 | 0.105 | 1.55 | 5890.60 |

Table S2. Primers used for amplification and cloning of TIPs and TSWV genes into pENTR-D/TOPO.

| Name | Sequence (5'-3') |
| --- | --- |
| ENTR-cyclophilinF | CACCATGGCACTTGCGCTCAGGTC |
| ENTR-cyclophilinR | TGACAGCTGGCCACAGTCAG |
| ENTR-enolaseF | CACCATGCCTGTCAAGTCCGTCAA |
| ENTR-enolaseR | GACAGGTTTGCGGAAGTTCT |
| ENTR-CP-VF | CACCATGCTGCCGTCGTTAGCCATCGTC |
| ENTR-CP-VR | GTATCGAGGGTCGTAGGGGG |
| ENTR-endoCP-G_N_F | CACCATGAAATTCCTGATCATCGC |
| ENTR-endoCP-G_N_R | GAACTGGCTGACGAGGTTGC |
| ENTR-endoCP-VF | CACCATGAAGCTGGAACTGGTCGC |
| ENTR-endoCP-VR | GTAGCGCCGCTGGAAGGGCC |
| ENTR-TSWV-NF | CACCATGTCTAAGGTTAAGCTCACTAAGG |
| ENTR-TSWV-NR | AGCAAGTTCTGTGAGTTTTGCCTG |
| ENTR-TSWV-G_N_F | CACCATGAGAATTCTAAAACTACTAGAACTAGTCG |
| ENTR-TSWV-G_N_R | CATAGACATGGGCATTTGAGACAAAATGATC |
| ENTR-TSWV-G_N_R1353 | ACCTATTAAGATTTCAGTCACAAA |
| ENTR-TSWV-G_N_SR | AATGCTTTTTGAATATTTGATTATGCAATCTCTAAC |
| ENTR-TSWV-G_C_F | CACCATGTTGATCATTTTGTCTCAAATGCCC |
| ENTR-TSWV-G_C_R | AAGCCACCTATGGATTTCTCTCACCTTGTC |
| ENTR-TSWV-G_C_SR | ATAGCTTGCAATGAAATTGAATGGGC |

Table S3. Primers used for amplification and cloning of TIPs and TSWV genes into MbY2H vectors, pPR3N and pBT3-SUC.

| Name | Sequence (5'-3') |
| --- | --- |
| 3NcyclophilinSfiF | ATTAACAAGGCCATTACGGCCATGGCACTTGCGCTCAGGTCGAT |
| 3NcyclophilinSfiR | AACTGATTGGCCGAGGCGGCCGTTATGACAGCTGGCCACAGTCAGCAAC |
| 3NenolaseSfiF | ATTAACAAGGCCATTACGGCCATGCCTGTCAAGTCCGTCAAGGCCCGC |
| 3NenolaseSfiR | AACTGATTGGCCGAGGCGGCCGTTAGACAGGTTTGCGGAAGTTCTTG |
| 3NCP-VSfiF | ATTAACAAGGCCATTACGGCCATGCTGCCGTCGTTAGCCATC |
| 3NCP-VSfiR | AACTGATTGGCCGAGGCGGCCGTTAGTATCGAGGGTCGTAGGGGGTGGA |
| 3NendoCP-G_N_SfiF | ATTAACAAGGCCATTACGGCCATGAAATTCCTGATCATCGCTGCCCTC |
| 3NendoCP-G_N_SfiR | AACTGATTGGCCGAGGCGGCCGTTAGAACTGGCTGACGAGGTTGC |
| 3NendoCP-VSfiF | ATTAACAAGGCCATTACGGCCATGAAGCTGGAACTGGTCGCCCTGTGC |
| 3NendoCP-VSfiR | AACTGATTGGCCGAGGCGGCCGTTAGTAGCGCCGCTGGAAGGGCC |
| 3NmATPaseSfiF | ATTAACAAGGCCATTACGGCCATGGCTCTCCTCTCCGTTCGTCTCGCT |
| 3NmATPaseSfiR | AACTGATTGGCCGAGGCGGCCGTTATTTACTTGCTGCGTTGAAACTGG |
| pBT3-G_N_SfiF | ATTAACAAGGCCATTACGGCCGTAGAGATAATTCGTGGAGACCATCCT |
| pBT3-G_N_SfiR | AACTGATTGGCCGAGGCGGCCCCACCTATTAAGATTTCAGTCACAAA |
| 3NendoCP-G_N_176SfiR | AACTGATTGGCCGAGGCGGCCGTTACTTGTACTTGGGCTGGGCGGAGG |
| 3NendoCP-G_N_189SfiR | AACTGATTGGCCGAGGCGGCCGTTAGATGCCGGCGGGGTCGAACTCCTG |
| 3NendoCP-G_N_177SfiF | ATTAACAAGGCCATTACGGCCATCCTCAGCCAGGTTCAGGAGTTCGA |
| 3NendoCP-G_N_190SfiF | ATTAACAAGGCCATTACGGCCTACCGTGTGAACTTCCAGACCGAGAA |
| pBT3-NSfiF | ATTAACAAGGCCATTACGGCCATGTCTAAGGTTAAGCTCAC |
| pBT3-NSfiR | CTGATTGGCCGAGGCGGCCTTAGCAAGTTCTGTGAGTTTTGCCTG |

Table S4, Number of dissected *F. occidentalis* first instar larvae that were immunolabeled with specific antisera against each TIP and was visualized by confocal microscopy from two replicates. MG, midgut; TSG, tubular salivary gland; PSG, principle salivary gland.

| Antisera | Number of dissected intact MG | Number of dissected intact TSG | Number of dissected intact PSG |
| --- | --- | --- | --- |
| No antibody control | 64 | 22 | 25 |
| Cyclophilin pre-immune serum | 47 | 10 | 21 |
| Enolase pre-immune serum | 29 | 13 | 10 |
| CP-V pre-immune serum | 38 | 13 | 18 |
| EndoCP-G_N_ pre-immune serum | 11 | 2 | 7 |
| EndoCP-V pre-immune serum | 28 | 12 | 10 |
| Anti-cyclophilin | 26 | 15 | 10 |
| Anti-enolase | 58 | 25 | 21 |
| Anti-CP-V | 35 | 6 | 11 |
| Anti-endoCP-G_N_ | 91 | 32 | 35 |
| Anti-endoCP-V | 50 | 20 | 20 |
| Anti-mATPase | 35 | 16 | 25 |
