## Supplementary figures and images for "Discovery of novel thrips vector proteins that bind to the viral attachment protein of the plant bunyavirus, tomato spotted wilt virus"

### Fig S1

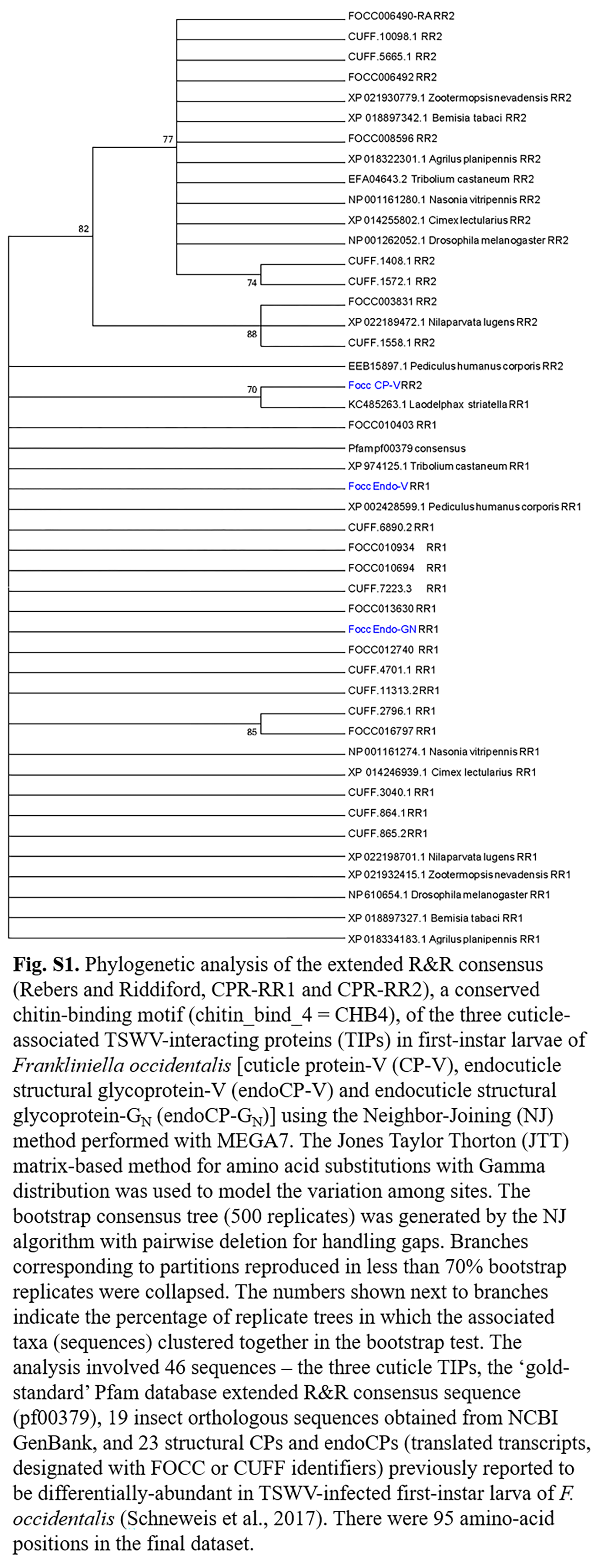

### Fig S2

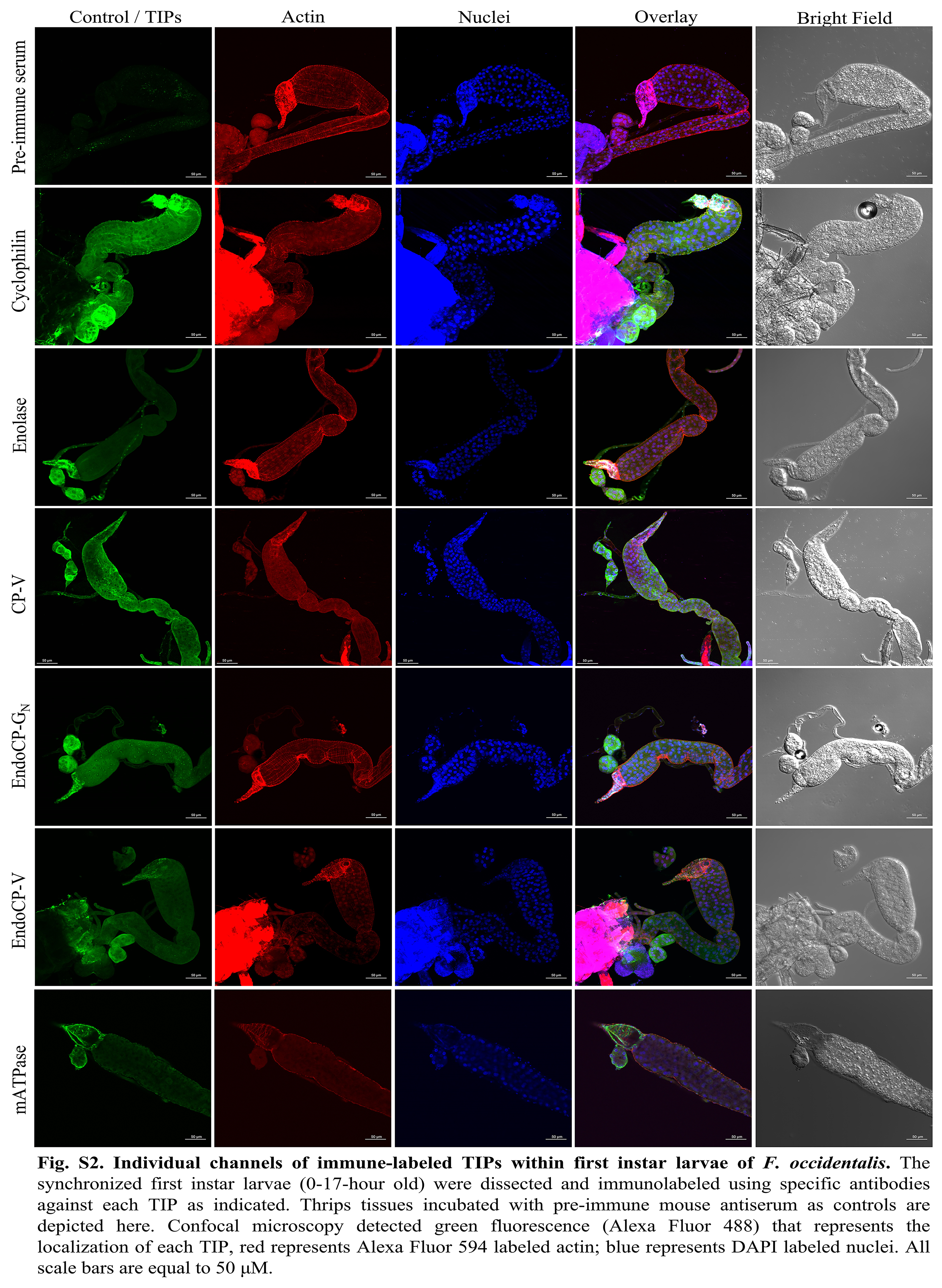

### Fig S3

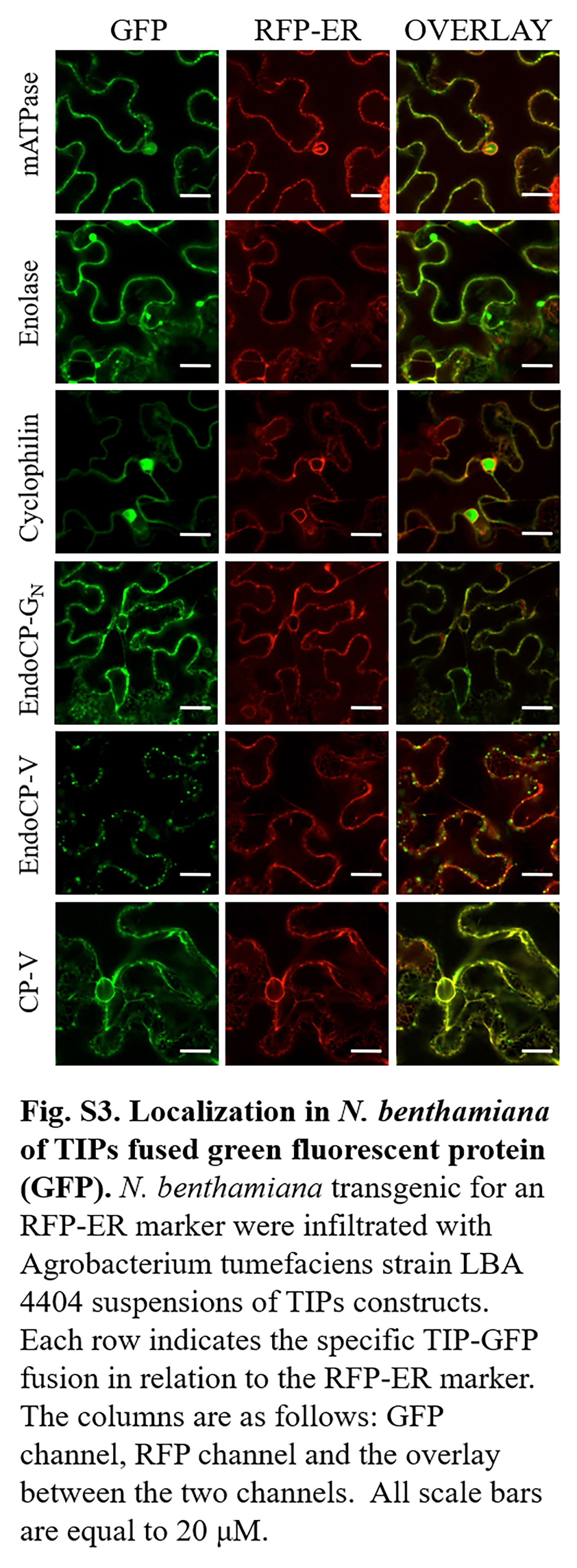

### Fig S4

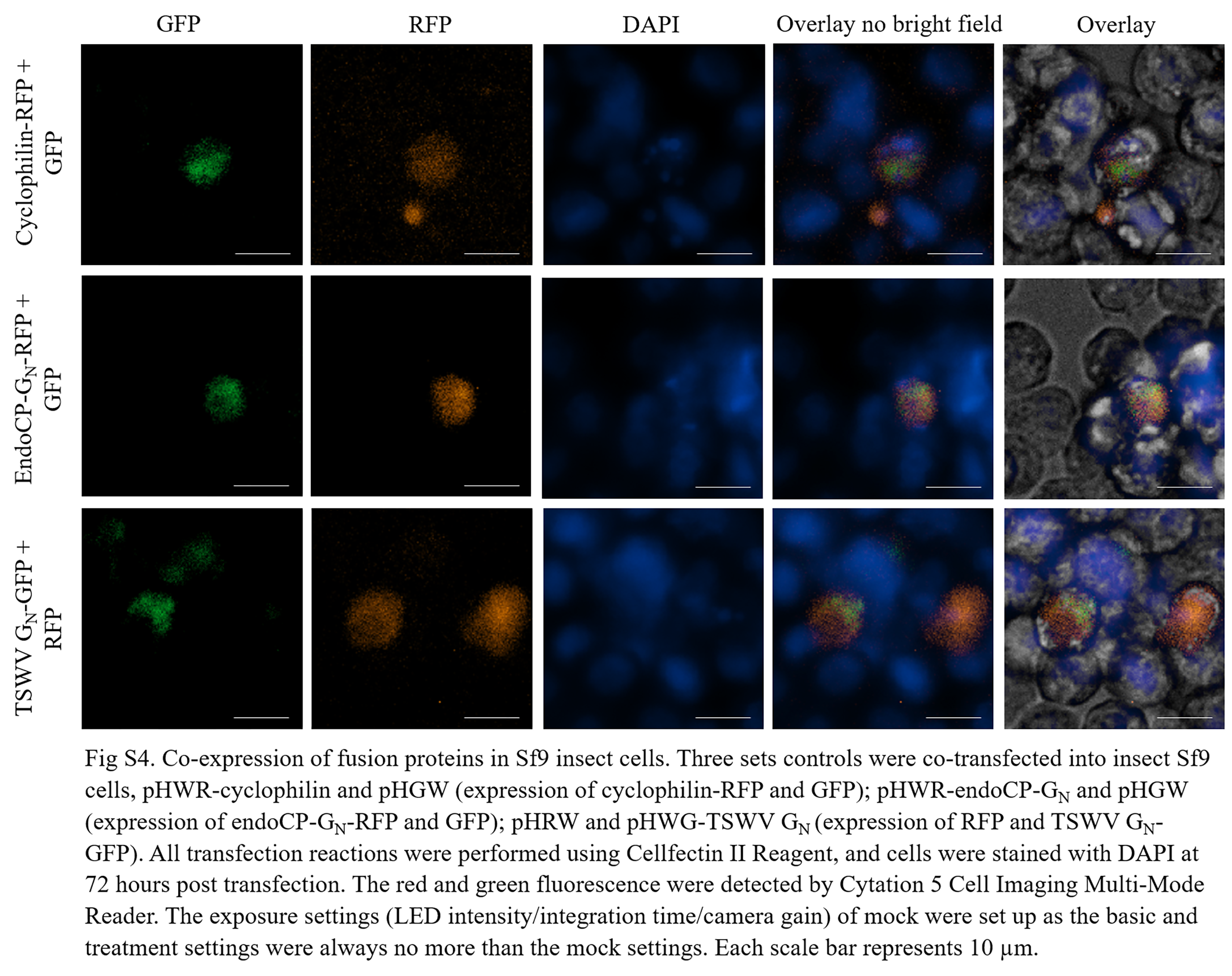
